# Variation and selection at predicted G-quadruplexes across the human pangenome

**DOI:** 10.64898/2026.06.18.733261

**Authors:** Saswat K. Mohanty, Maximillian G. Marin, Linnéa Smeds, Francesca Chiaromonte, Christian D. Huber, Kateryna D. Makova, Human Pangenome Reference Consortium

## Abstract

G-quadruplexes (G4s), non-canonical DNA structures whose sequence motifs occupy approximately 1% of the human genome, are important for myriad cellular functions, including regulating transcription and replication. Yet they also contribute to genomic instability by increasing mutations and structural variation. Despite their significance, G4 motifs have not been studied in detail across multiple human genomes. Here, we conducted a comprehensive analysis of presence/absence and sequence variation, measured selection strength, and evaluated gene expression regulation potential for predicted G4s (pG4s) across population groups in the second release of the Human Pangenome Reference Consortium dataset, comprising high-quality, near-telomere-to-telomere diploid genomes from 231 individuals worldwide, along with three reference assemblies. Across the human pangenome, we identified over 353 million pG4s, including 1.15 million pG4s absent from reference assemblies but shared across other haplotypes. Our analysis revealed that pG4 sharing patterns recapitulate human population structure: African individuals displayed lower levels of pG4 sharing than non-Africans, whereas East Asian individuals exhibited higher levels of sharing. By analyzing the site frequency spectrum across various genomic annotations, we computed and compared selection coefficients (*S_d_*) at pG4 vs. non-pG4 sites. As expected, the strongest purifying selection (*S_d_ ≥ 10*) was detected at protein-coding exons, where pG4 sites had similar or lower selection coefficients compared with those for pG4 sites. Strikingly, this pattern reversed at regulatory regions: although purifying selection was weaker overall at promoters, introns, enhancers, and replication origins (*1 ≤ S_d_ < 10*), pG4 sites at these regions experienced stronger selection than non-pG4 sites—suggesting that pG4s play functional roles outside coding sequences. Additionally, by integrating pG4 data with long-read transcriptome data profiles from this large cohort, we found that pG4s located at promoters and at (or near) exon-intron junctions may influence variation in gene expression levels and transcript isoforms, respectively, across the human pangenome individuals. Leveraging extensive population-scale data, our research illuminates the fundamental importance and functional relevance of G4s across human genomes.

## INTRODUCTION

Whereas the majority of the human genome folds into the right-handed B-DNA double helix (Watson and Crick 1953), certain segments can transition into various non-B DNA conformations (Kouzine et al. 2017). These alternative structures, which include Z-DNA (Mitsui et al. 1970), cruciforms (Panayotatos and Fontaine 1987), H-DNA (Felsenfeld and Rich 1957), bent DNA (Prosseda et al. 2004), and G-quadruplexes (G4s) (Sen and Gilbert 1988), are transient and less stable than B DNA (Varizhuk et al. 2019), yet they play important biological roles in replication, transcription, chromatin organization, and recombination, as well as in other cellular processes (reviewed in (Wang and Vasquez 2022)).

G4s form via Hoogsteen (Hoogsteen 1963) hydrogen bonds between non-sequential guanines, which create planar G-tetrads that are further stabilized by monovalent cations such as potassium (Sokulska et al. 2026). Structurally, a G4 consists of stems—composed of two to five consecutive guanines—and loops of one to twelve nucleotides (Guédin et al. 2010). These structures may also feature "bulges", where the guanine runs in the stems are interrupted by other nucleotides (Mukundan and Phan 2013; Papp et al. 2023) (Fig. 1). G4s can be found in both DNA and RNA (Fay et al. 2017) and can manifest as either intra- or intermolecular structures (Zhang et al. 2025).

**Figure 1.**
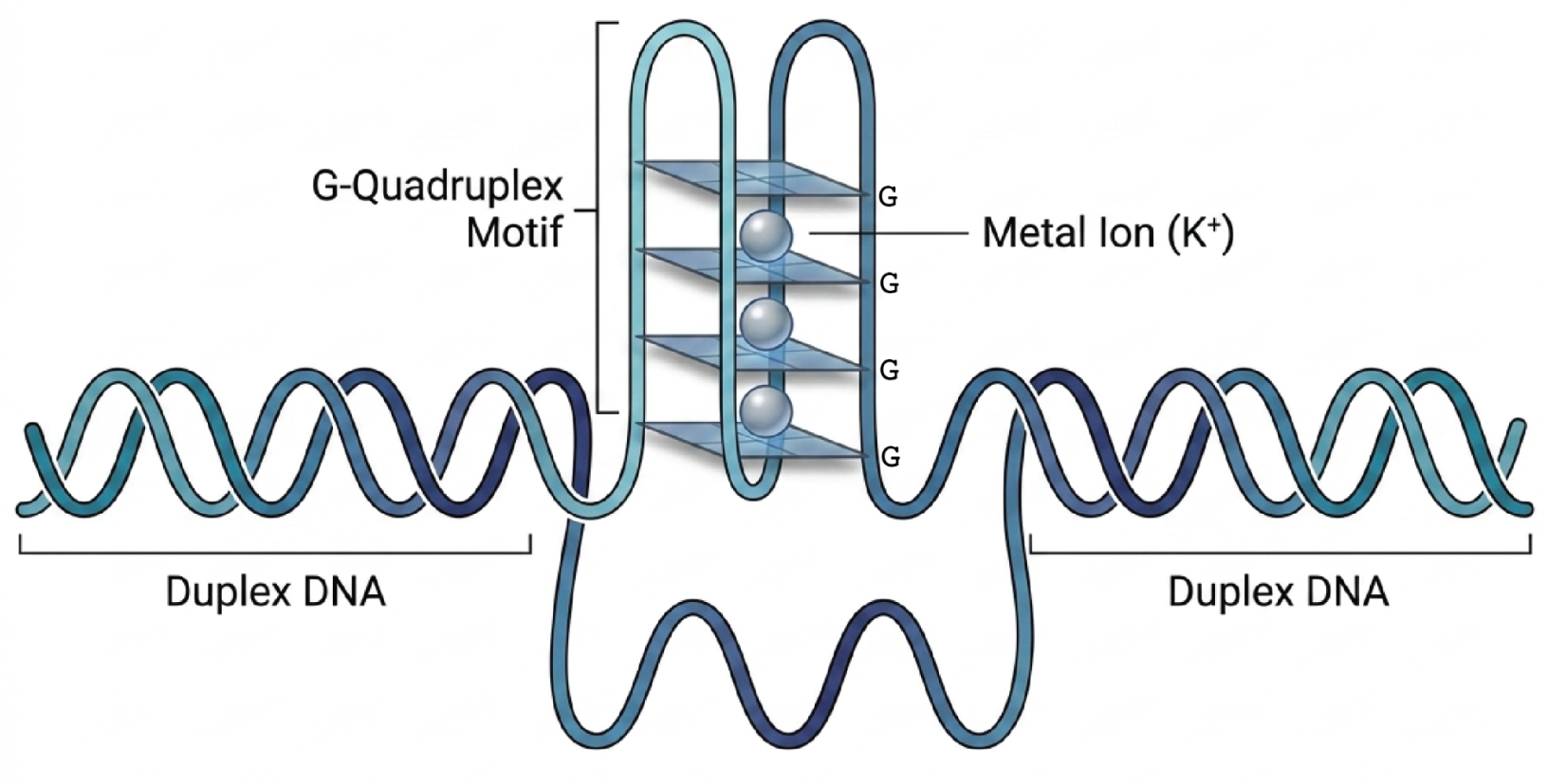
A G-rich motif showing the G-quadruplex structure with potassium ions between the tetrads. Figure adapted from (G. Wang and Vasquez 2022).

G4s are hallmarks of genomic instability (Teng et al. 2021; Guiblet, Cremona, et al. 2021; Ravichandran et al. 2019; Zhang et al. 2024; Pavlova et al. 2021). They have been identified as potential mutation hotspots for germline (Guiblet et al. 2018; Guiblet, Cremona, et al. 2021) and somatic mutations, including large-scale genomic rearrangements, during cancer (Georgakopoulos-Soares et al. 2018; Stein and Eckert 2021; Zhang et al. 2023). In the context of DNA replication, G4s appear to serve dual, conflicting roles. On the one hand, they can act as obstacles that stall replication forks (Stein et al. 2022), thereby fostering genomic instability (Sun and Hurley 2010; Lormand et al. 2013). On the other hand, they likely facilitate the firing of replication origins, where they are overrepresented and their absence can hinder replication initiation (Prioleau 2017; Valton et al. 2014). In addition, G4s also fulfill other important functions (reviewed in (Wang and Vasquez 2022)). For instance, these structures are particularly prevalent at telomeres, where they participate in telomere maintenance (Bryan 2020; Lin and Yang 2017; Moye et al. 2015; Xu and Komiyama 2023). Moreover, they have been shown to be enriched at immunoglobulin switch regions (Vallur and Maizels 2008) and thus may contribute to class-switch recombination, i.e., an essential biological process in which activated B cells change the class of antibody they produce.

G4s’ influence on transcriptional regulation is multifaceted. First, G4s are significantly enriched at cis-regulatory elements, particularly at promoters and enhancers, where they serve as recruitment sites for transcription factors (Spiegel et al. 2021), often with higher binding affinities than B-DNA motifs (Zhang et al. 2024) and thus boost expression (C.-Y. Lee et al. 2020). Indeed, a recent study showed that inserting a G4 into the promoter region of a synthetic reporter gene enhanced its transcription (Chen et al. 2024). A handful of studies examined G4-mediated transcriptional regulation at particular genes, including *c-MYC* (Esain-Garcia et al. 2024), *BCL-2* (Wang et al. 2010), *KRAS* (Cogoi and Xodo 2006), *HRAS* (Cogoi et al. 2014), *MET* (Yan et al. 2016), *PDGFRn* (Brown et al. 2017), *AKR1C* (Yuan et al. 2023), *IRF8*, and *VEGFA* (Gong et al. 2021). Second, G4s may impede RNA polymerase progression and arrest transcription (Broxson et al. 2011; Smestad and Maher 2015). Third, G4s are instrumental in shaping three-dimensional chromatin architecture and facilitating RNA polymerase II-mediated long-range interactions, such as enhancer-promoter loops that actively upregulate gene expression (Yuan et al. 2023). G4s’ strong association with open chromatin (Spiegel et al. 2021; Lago et al. 2021) and active epigenetic marks (such as H3K4me3) further underscores their role in regulating transcriptional activity (Zhang et al. 2024).

Fourth, G4s have been identified as regulators of alternative splicing (Varshney et al. 2020; Georgakopoulos-Soares, Parada, Wong, et al. 2022; Georgakopoulos-Soares, Parada, and Hemberg 2022; Peynet et al. 2026). A recent study (Georgakopoulos-Soares, Parada, Wong, et al. 2022) indicated that G4s can occur on either template or non-template strand, but are preferentially located on the non-template strand, with their frequency reaching a peak approximately 50 base pairs from the splice site junction within introns (Georgakopoulos-Soares, Parada, Wong, et al. 2022). It was also shown that the formation of a G4 on the non-template strand, when it is expressed as an RNA G4, e.g., at intron 3 of *TP53* (Marcel et al. 2011), intron 1 of *PAX9* (Ribeiro et al. 2015), and intron 15 of *IGFN1* (Verma and Das 2018), might trigger alternative splicing as a direct effect. At the same time, it has also been discussed that G4s present on the template strand, may lead to a slowdown of RNA polymerase II, which might induce the exposure of weak splice sites, altering the exon definition as an indirect effect (Peynet et al. 2026; Georgakopoulos-Soares, Parada, and Hemberg 2022). Validating this, a prior multi-species study demonstrated that, in humans, template G4s are significantly associated with exon skipping as compared to non-template G4s (Tsai et al. 2014). However, to date, the lack of comprehensive, genome-wide G4 studies has restricted these gene regulatory investigations to a limited set of candidate genes, leaving the broader landscape of G4-driven regulation largely unexplored, particularly in the context of gene expression variation in human populations.

Consistent with these essential regulatory mechanisms, some G4 motifs have been shown to evolve under purifying selection across various functional genomic regions (Guiblet, DeGiorgio, et al. 2021). In addition, the central guanines of the 3G-tracts in canonical (i.e., bulge-less) G4s located at promoters were found to be under strong selection pressure—comparable to that for missense mutations in protein-coding genes (Li et al. 2023). Similarly, a recent study in great ape telomere-to-telomere (T2T) genomes highlighted that G4s’ stems experience stronger purifying selection as compared to loops (Zhang et al. 2026) when located within promoters, enhancers, and 5’ UTRs (untranslated regions). Despite the importance of G4s’ regulatory roles, the selection coefficients at these loci have not been quantified. It is currently unclear whether G4s situated in regulatory regions beyond promoters are subject to selective constraints similar to those observed in protein-coding exons. Furthermore, because G4s are integral components of different genic compartments (Guiblet, DeGiorgio, et al. 2021), the selective pressure acting on these regions with respect to the regions surrounding them, while accounting for different mutation rates between G4 and non-G4 sites (Guiblet, Cremona, et al. 2021; Du et al. 2014), remains to be quantified.

Multiple population-level analyses have uncovered important patterns of polymorphism at G4 loci across the genome. An initial investigation using 1000 Genomes Project data concluded that more than 96% of promoter G4 motifs lacked polymorphisms, and that single-nucleotide polymorphisms (SNPs) disrupting these structures altered promoter activity (Baral et al. 2012). A subsequent study (Gong et al. 2021) expanded on this by characterizing single-nucleotide variants (SNVs) at G4s highlighting millions of gains, losses, or structural variants at predicted G4s, most of them near transcription start sites (TSS), that can substantially affect the expression of host genes. More recently, a pangenome-scale analysis (Chantzi et al. 2025) employing the initial Human Pangenome Reference Consortium data (i.e., HPRCv1) (Liao et al. 2023) confirmed that G4s contribute to genomic instability, evidenced by an elevated frequency of all mutation types at their motifs. However, this first pangenome release included only 47 individuals and covered only ∼91% of common variants (allele frequency ≥1%) and 62% of rare and common variants (allele frequency ≥0.1%) (Human Pangenome Reference Consortium 2026). Therefore, a larger sample size of high-quality sequences is required to increase power and to uncover a more comprehensive picture of variation at G4s in human genomes.

The availability of high-quality T2T reference genomes has facilitated comprehensive studies of previously inaccessible complex genomic regions (Makova et al. 2026). These T2T assemblies, including long sequencing reads with minimal error rates at non-B DNA sites (Makova and Weissensteiner 2023), provide more accurate sequence information at these loci (Rhie et al. 2023; Smeds et al. 2025). Such information, combined across hundreds of human genomes, offers an unprecedented opportunity to study the complete repertoire of G4s and selection acting on them.

In this study, we used the second release of the HPRC (HPRCv2), comprising 232 diploid human genomes from populations worldwide (Human Pangenome Reference Consortium 2026). These genomes were generated using a combination of short- and long-read technologies and assembled with state-of-the-art algorithms, e.g., hifiasm (Cheng et al. 2022, 2021). Utilizing HPRCv2, which covers more than 99% of all common variants and about 85% of all rare and common variants, we addressed the following questions. First, does variation in G4s’ predicted presence/absence, and variation in their sequences, reflect human population structure? Second, how strong is selection acting on sites within and outside of G4s in different functional genomic regions, and what does it tell us about the functional potential of these non-B DNA loci? Third, what does integrating long-read expression data unveil about the effect of G4s’ genetic variation at promoters and exon/intron junctions on RNA expression across haplotypes and individuals? Taken together, using a pangenome approach, we link pG4s to human population structure, regulatory roles, and selection, establishing G4s as functionally important elements of the human genome.

## RESULTS

### G4 prediction and characterization across the human pangenome

We considered the 234 human genomes (466 haplotypes) included in HPRCv2 (Human Pangenome Reference Consortium 2026). This dataset consisted of 231 haplotype-resolved, near-T2T genomes from diploid individuals, along with three reference assemblies: the diploid, haplotype-resolved HG002, and the haploid CHM13 and GRCh38. A total of 118 of the 234 genomes included the Y chromosome.

We used G4Discovery (Mohanty et al. 2025) to detect G4 motifs in each of these 466 haplotypes, identifying a total of ∼353 million predicted G4s (later called “pG4s”) across the pangenome, with an average of 7.58×10^5^ pG4s per haplotype (Table 1; see Methods). Excluding sex chromosomes, the number of pG4s varied across human haplotypes with an approximately normal distribution centered at 7.27×10^5^ pG4s (Fig. 2A). The number of pG4s on the X chromosome also showed a bell-shaped distribution centered at 3.2×10^4^ pG4s per haplotype. Most Y chromosomes had between 5×10^3^ and 7×10^3^ pG4s per haplotype.

**Figure 2.**
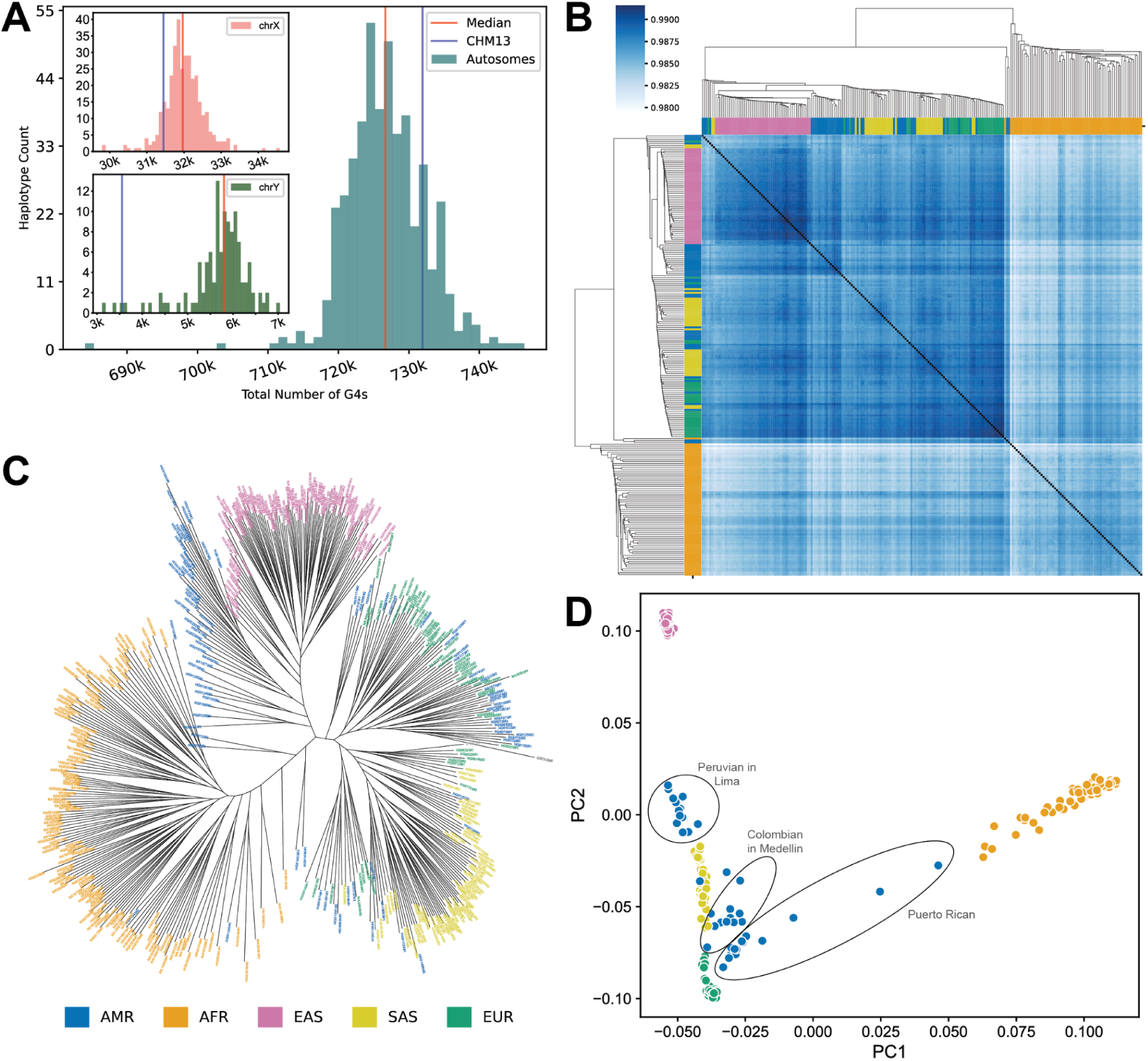
Population stratification using population-level pG4 data. **A**. Histogram showing distribution of the total number of pG4s across autosomes and sex chromosomes. **B**. Heatmap showing the proportion of pG4s shared between any two haplotypes. Superpopulation color labels are shown in panel C. **C**. Unrooted maximum-parsimony phylogenetic tree, derived using the pG4 presence-absence variation matrix, showing superpopulation stratification. **D**. PCA plot using the pG4 SNP data referenced on CHM13. The plot utilizes five superpopulation labels as annotated in C.

**Table 1.** The number of pG4s annotated in the HPRCv2 haplotypes. The number of pG4s annotated using G4Discovery (Mohanty et al. 2025) across all 466 human haplotypes comprising 231 diploid human assemblies and three reference assemblies: CHM13 (1), GRCh38 (1) and HG002 (2). The median number of pG4s per haplotype was calculated by dividing the total number of pG4s by 466. The number of pG4s analyzed in the presence-absence data is also presented.

| **pG4 counts** | **Total N of pG4s across all 466 haplotypes** | **Median N of pG4s per haplotype** |
| --- | --- | --- |
| pG4s across the pangenome | 353,549,535 | 762,787 |
| pG4s at sex chromosomes (X & Y) | 11,870,954 | - |
| pG4s removed from analysis due to unassigned or flagged regions | 22,585,609 | 47,737 |
| pG4s absent from the pangenome alignment (autosomes only) | 2,240,186 | 4,231 |
| pG4s retained for the analysis | 317,914,256 | 682,159 |
| **Presence-absence pG4 variation (autosomes only)** | **1,095,105 pG4 families** |  |
| pG4s in fixed pG4 families | 251,901,687<br>(506,236 families) | 540,544 |
| pG4s in variable pG4 families* | 65,650,932<br>(228,876 families) | 140,900 |
| pG4s in haplotype-specific pG4 families | 361,637<br>(359,993 families) | 715 |

\*For variable pG4s families, some haplotypes (more than one) had a pG4 while others did not have a pG4 at an orthologous location.

### pG4 presence/absence and sequence variation reveal human superpopulation structure

We next examined the behavior of pG4s as population genetics markers. We excluded pG4s (1) located on sex chromosomes, in flagged regions of low assembly confidence (see Methods), or in unassigned scaffolds (an average of 47,737 pG4s per haplotype, or ∼6.39% from the total dataset), and (2) absent from the pangenome alignments (an average of 4,231 pG4s per haplotype, or ∼0.6% from the total dataset). We constructed a presence/absence matrix for the remaining pG4s across the 466 haplotypes. In this matrix, a pG4 present at a given orthologous position in some haplotypes could be absent in others due to: (1) mutations that cause the sequence to no longer be predicted as a pG4, or (2) deletion of the region harboring the pG4. We used this matrix to analyze the population- and superpopulation-level structure of the human pangenome.

We then defined “pG4 families” as collections of pG4s at orthologous locations across human haplotypes. We categorized these pG4 families into fixed (shared across all haplotypes), variable (shared across multiple haplotypes but not fixed), and haplotype-specific (pG4s unique to a single haplotype). Consistent with the high degree of similarity expected across human genomes, fixed pG4 families comprised approximately 46% of all families but accounted for over 79% of all pG4s in our analysis. Variable pG4 families comprised 21% of all families and contained approximately 21% of all pG4s. In contrast, haplotype-specific families (found mostly in African and admixed American populations; Fig. S1) comprised 33% of all families but contained only ∼0.1% of all pG4s. The number of haplotype-specific pG4s ranged from as few as 43 (HG002, haplotype 2) to as many as 2,494 (HG02965, haplotype 1) per haplotype (Fig. S2). Notably, we found as many as 1,150,535 pG4s (∼2,490 pG4s per haplotype) absent from the reference assemblies (CHM13, GRCh38, and HG002) but shared across several haplotypes in the pangenome (Fig. S3).

To assess the relatedness of haplotypes based on their pG4 sharing, we calculated the proportion of pG4 families shared between any two haplotypes and applied cluster analysis (heatmap in Fig. 2B; see Methods). African haplotypes exhibited lower sharing than non-African haplotypes, while East Asian haplotypes clustered tightly together. The patterns of population-level sharing were broadly similar to those observed at the haplotype level (Fig. S4). Therefore, clustering based on pG4 sharing displays the relationships among human superpopulations and populations.

Next, we used the pG4 presence/absence matrix to reconstruct the most parsimonious phylogenetic tree for the 466 HPRCv2 haplotypes (Fig. 2C). Color-coding tree branches by superpopulation, we observed haplotypes forming distinct clades for the African and East Asian superpopulations (Fig. 2C). In contrast, color-coding tree branches by specific population, most groups did not resolve into distinct lineages, likely reflecting recent or more ancient admixture (Fig. S5). An exception to this was the population of Peruvians (PEL), in which most haplotypes formed a single clade. Thus, both the clustering and the phylogenetic analysis, based on pG4 families sharing support: (1) a distinct genetic composition of superpopulations, with less structured differentiation among populations; (2) an out-of-Africa dispersal pattern, with African populations being the most diverse; and (3) an interspersed pattern of admixed American haplotypes (except for Peruvians) across clusters and the phylogenetic tree.

We also computed the number of population-specific pG4s, defined as pG4 families shared by at least two haplotypes of the same population but absent from the other populations. We found that African populations (e.g., the Gambian population, GWD) and some recently admixed population groups (e.g., PEL and Puerto Ricans, PUR) had a particularly high number of population-specific pG4s (Fig. S6).

Finally, we investigated the grouping of the 466 haplotypes based on sequence-level variation within pG4 loci genome-wide. We performed a Principal Component Analysis (PCA) using PLINK2 (Chang et al. 2015) (Fig. 2D; see Methods). The first principal component clearly separated African haplotypes into a distinct group. The second principal component separated East Asian, South Asian, and European superpopulations. In contrast, haplotypes from populations with recent admixture—specifically PEL and those from Puerto Rico, Colombia (CLM)—showed higher levels of intra-population variance and were located near (for PEL) or interspersed among (for CLM and PUR) the European and South Asian haplotypes (Fig. 2D). Altogether, our results suggest that both the presence/absence of pG4s and their sequence variation reflect the demographic history of superpopulations and populations in the human pangenome.

### Purifying selection acting on pG4s

Using sequence-level variation data and considering different types of functional genomic regions separately, we quantified the selection pressure acting on genetic variation within and outside of the pG4 sites. We restricted our analysis to four of the 27 HPRCv2 populations, which were selected based on the following criteria: (1) they had a large enough number (*N*≥13) of individuals to infer the distribution of fitness effects (DFE), and (2) individuals belonging to these populations clustered closely with each other in the PCA analysis (Fig. S7). The populations meeting these criteria were Gambian in Western Division Ð Mandinka (GWD; *N*=17), Japanese in Tokyo (JPT; *N*=16), Peruvian in Lima (PEL; *N*=15), and Kinh in Ho Chi Minh City (KHV; *N*= 13). For these populations, we computed and contrasted selection coefficients within (“pG4”) and outside (“non-pG4”) pG4s located in promoters, 5’ UTRs, protein-coding sequences (CDS), introns, 3’ UTRs, enhancers, and origins of replication (see Methods). For each of these regions, we inferred ancestral allele states for the biallelic SNPs using chimpanzee and orangutan as outgroups, and constructed unfolded site-frequency spectra (see Methods). After this, fitness effects were inferred using fastDFE (Sendrowski and Bataillon 2024), following downsampling of all populations to 25 haplotypes via hypergeometric projection (see Methods). Non-genic non-repetitive (NGNR) regions were used as a neutral background for this calculation (see Methods). Specifically, the inference concerned the parameters of a gamma-distributed DFE, with selection effects expressed in terms of the population-scaled selection coefficient, *S_d_* = 4*N_e_s*, where *N_e_* is the effective population size, and *s* is the selection coefficient. Thus, the distributions of selection coefficients could be compared across different functional regions for the same population. Additionally, we compared the rankings of average selection coefficients measured between pG4 and non-pG4 sites at different functional regions across the four populations under analysis. This quantifies the distinct selection pressure on pG4 versus non-pG4 sequences, aiding in assessing selection acting directly on the pG4s themselves rather than on their genomic environment.

We made three main observations. First, consistent with expectations, we observed high average selection coefficients for CDS across all four populations (Fig. 3A-D), validating our approach, as strong purifying selection is expected at coding sequences. At CDS regions, we further separately considered 0-fold degenerate sites (where any possible nucleotide change alters the amino acid) and 4-fold degenerate sites (where changing the nucleotide for any of the other nucleotide results in the same amino acid, and which are frequently used as a neutral baseline), in order to quantify differences between purely nonsynonymous and purely synonymous mutations, respectively. We found that selection coefficients at 0-fold-degenerate sites were among the highest in our overall analysis—with average *S_d_* values around 40 or higher, consistent with strong purifying selection. The model’s inference of selection coefficients for 4-fold degenerate sites was generally underconfident due to the limited number of sites. However, we still observed lower selection coefficients at 4-fold degenerate sites than at 0-fold degenerate sites, as predicted (Fig. 3A-D). Notably, the strength of selection at CDS remained similar regardless of whether the sites were within or outside pG4s, or was higher for non-pG4 sites.

**Figure 3.**
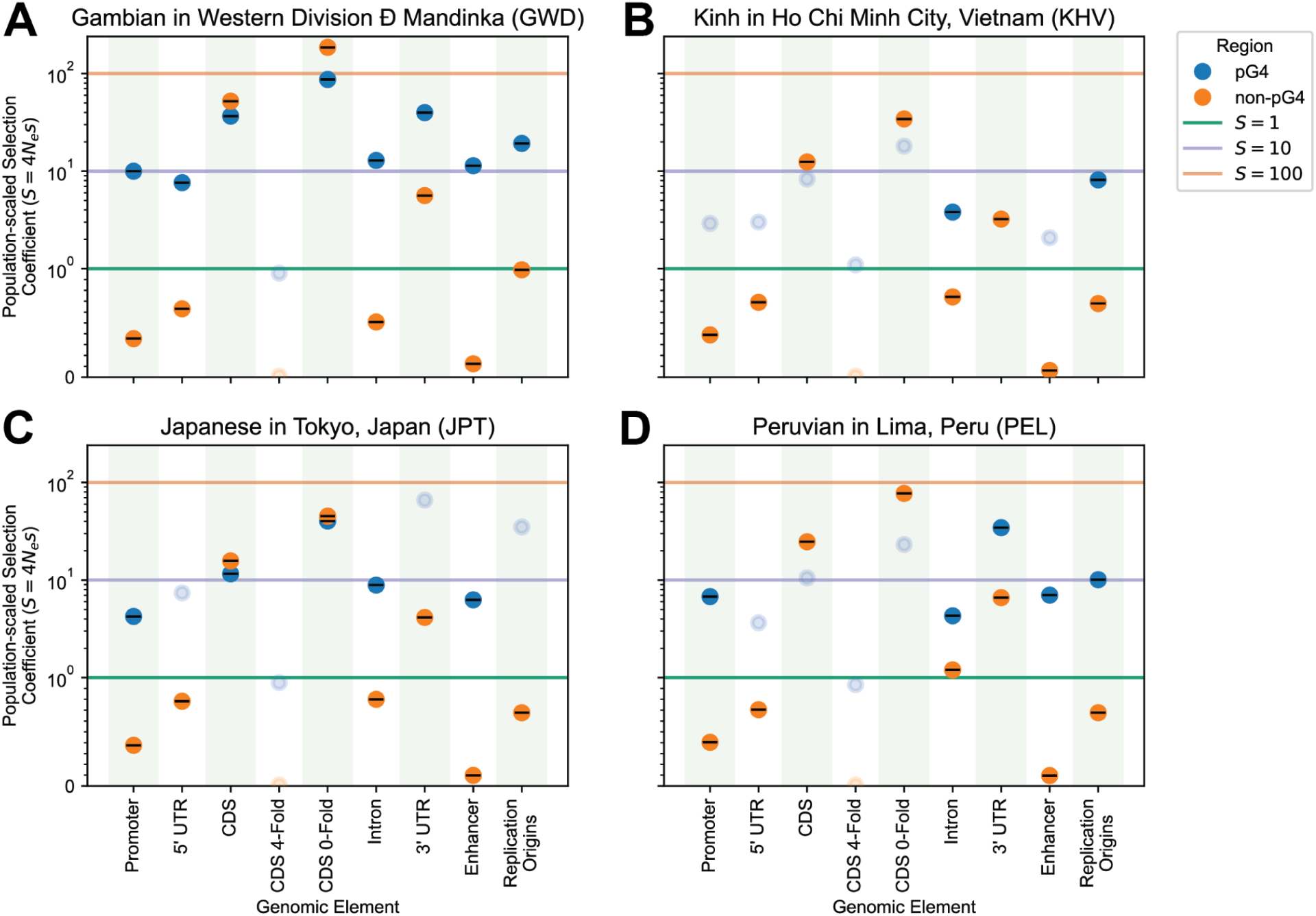
Population-scaled selection coefficients at in-pG4 and non-pG4 loci, across different functional genomic regions. The dot plot shows the population-scaled selection coefficient (*S* = 4*N_e_s*) for four different populations: **A**. Gambian in Western Division Ð Mandinka (GWD), **B**. Japanese in Tokyo (JPT), **C**. Peruvian in Lima (PEL), and **D**. Kinh in Ho Chi Minh City (KHV). Functional genomic regions are shown on the x-axis. The horizontal lines drawn at *S* = 1 marks the boundary between neutral (*S_d_* < 1) and weakly deleterious selection (1 *≤ S_d_* < 10), at *S* = 10: between weakly and moderately deleterious selection (10 ≤ *S_d_* < 100), and at *S* = 100: between moderately and strongly deleterious selection (*S_d_* ≥ 100). Circles in light colors indicate regions where the model didn’t fit the observed data with high confidence.

Second, across several other functional regions examined, pG4s predominantly exhibited selection coefficients consistent with weak (*1 < S_d_ < 10*) to moderate/strong (*S_d_ ≥ 10*) levels of purifying selection (Fig. 3A-D). Notably, the pG4 intervals in promoters, introns, and enhancers remained within or at the boundary of the weakly or moderate/strong deleterious selection regime, suggesting an essential functional role for these pG4s. In contrast, non-pG4 sites in these regions appeared to evolve neutrally. The model estimated that pG4s within 5’ UTRs were subject to weak selection in the Gambian population, while pG4s within 3’ UTRs were subject to moderate-to-strong purifying selection in the Gambian and Peruvian populations; the other populations followed the same trend, however, for them the model was underconfident (likely due to the small number of pG4s; Fig. 3A-D). For transcribed regions, we detected evidence of stronger purifying selection for pG4s located on the template vs. non-template strand, particularly for introns (which possessed the largest number of sites allowing for a statistically meaningful analysis), but the difference was relatively small (Fig. S8). In summary, our results were consistent with weak-to-moderate/strong purifying selection for pG4s at promoters, introns, enhancers, and possibly also UTRs, whereas the surrounding non-pG4 sequences appear to evolve neutrally.

Third, our analysis provided evidence of purifying selection acting on pG4s at core replication origins across the four populations considered. As described previously (Akerman et al. 2020), *core* replication origins are high-activity, shared-across-tissues origins, from which 80% of replication initiation takes place, whereas *stochastic* origins are low-activity origins that are mostly cell-type-specific. Our findings indicate that variation at pG4s in core replication origins is under strong selective pressure, with coefficients indicating moderate to strong selection levels across all populations, whereas non-pG4 regions exhibited low selection coefficients consistent with neutral evolution. While model inference for *core* origins was largely stable across populations, it was largely unstable for the pG4 sites in *stochastic* origins (Fig. S9). Non-pG4s at *stochastic* origins also showed near-neutral selection coefficients, similar to such sites at core origins.

Overall, our results point to four different types of functional regions of the genome—promoters, enhancers, introns, and core replication origins—where pG4 sites experience stronger selection than surrounding sequences. Moreover, pG4 sites in these functional regions form the next tier of selection-coefficient strength—ranging from weak to moderate/strong—after CDS sites, where purifying selection is expected and was validated by our approach. pG4 sites in 5’ and 3’ UTRs also likely evolve under purifying selection—although in some populations, we lack statistical support for this claim. In contrast, evolution of non-pG4 sequences in most of these functional regions (all but CDS and 3’ UTRs) is consistent with neutrality.

### Genetic variation at pG4s proximal to TSSs is associated with differences in gene expression levels across the pangenome

We next tested a hypothesis that pG4 genetic variation at promoters or 5’ UTRs contributes to differences in gene expression levels among individuals in the human pangenome. We performed an expression quantitative trait loci (eQTL) analysis (see Methods) using the long-read Pacific Biosciences (PacBio) transcriptome data from lymphoblastoid cell lines (Human Pangenome Reference Consortium 2026; Marin and Li 2026) for 206 human pangenome samples (excluding the reference assemblies) as our phenotype. The eQTL analysis (using the window size of 20 kb; see Methods) yielded 4,868 significant variants across 992 genes (*q*-value < 0.01). Among the significant variants from this eQTL, we identified those that lie within 2 kb upstream and 1 kb downstream (a total of 3 kb) of the transcription start site (TSS) of the longest mRNA of protein-coding genes annotated in the human reference genome (CHM13). This resulted in 4,069 variants across 930 genes. A total of 239 of these variants overlapped with pG4s. Following this, we conservatively retained genes that had only pG4 variants significantly associated with their expression variation according to our eQTL and had no other significant variants. After filtering out pG4s on sex chromosomes and multi-mapping pG4 families, we were left with 16 candidate pG4 variants, one at each gene (Table S1).

Among these 16 pG4 variants (Fig. S10), three (in *POGLUT3*, *PLCB2*, and *MOCS2*) were predicted to substantially change G4 stability (as evidenced by pqsfinder (Hon et al. 2017) and G4Hunter (Bedrat et al. 2016) scores, Table 2) and four (in *MEGF9*, *SV2B*, *SAAL1*, and *DGCR6*) were predicted to disrupt G4 formation altogether (Table 2). To confirm the association of these variants with gene expression phenotypes, we compared normalized expression levels across the two homozygote classes and the heterozygotes at each locus (homozygotes for the less stable or disrupted pG4 allele were plotted as "0"). In six of seven cases (all but in *SV2B*), expression levels differed significantly between the two homozygote classes—that is, between individuals carrying two copies of the variant conferring a more vs. less stable G4, or between individuals carrying two copies of the variant that abolishes versus permits G4 folding. However, the effects of these variants on expression increase vs. decrease were different in each case. The predicted folding vs. nonfolding of a G4 was associated with increased gene expression levels for *DGCR6* and *SAAL1*, but with decreased levels for *MEGF9* (Fig. 4A-C). Similarly, more stable pG4 variants were associated with increased gene expression levels for *POGLUT3*, but decreased for *MOC52* and *PLCB2* (Fig. 4D-F).

**Figure 4.**
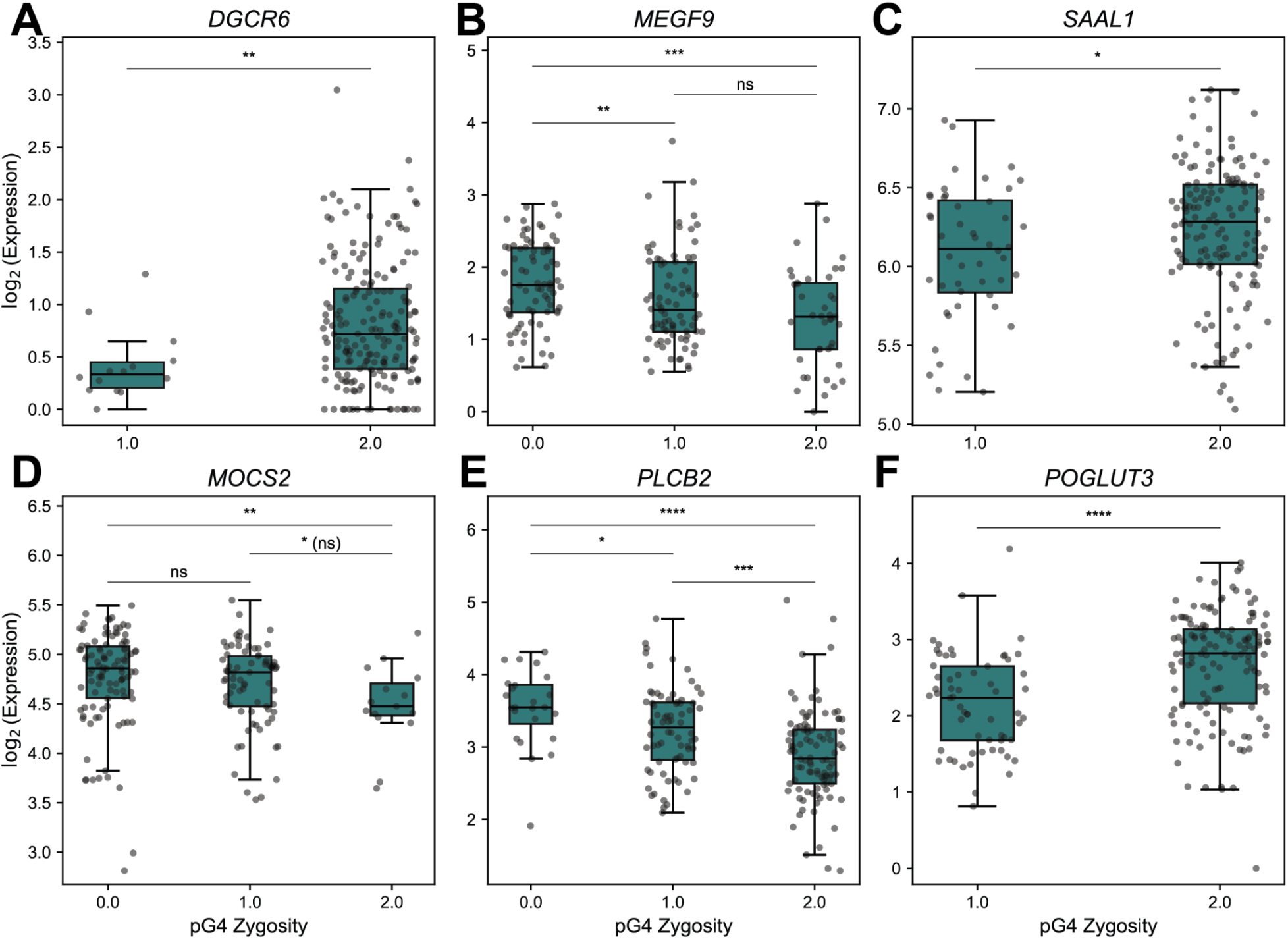
pG4-zygosity-dependent gene expression levels, across six candidate genes. The boxplot (with striplot over it) shows the log_2_-transformed counts per million (CPM) values versus the pG4 zygosity at six different candidate genes: (**A**) *DGCR6*, (**B**) *MEGF9*, (**C**) *SAAL1*, (**D**) *MOCS2*, (**E**) *PLCB2*, and (**F**) *POGLUT3*. Significance stars denote results from two-sided Mann-Whitney-Wilcoxon pairwise tests. (non-significant (ns): *p-value >* 0.05; *: 0.01 < *p-value* ≤ 0.05; **: 0.001 < *p-value* ≤ 0.01; ***: 1.00e-04 < *p-value* ≤ 0.001; ****: *p-value* ≤ 1.00e-04)

**Table 2.** Genes with a significant association between expression levels and genetic variants within pG4spG4s, as a result of eQTL analysis. This table lists candidate genes featuring a pG4 located proximal to the Transcription Start Site (TSS) that exhibit a strong positive or negative association with gene expression. For each gene, the orthologous pG4 coordinates within the CHM13 reference assembly are provided, along with the beta value and the *q*-value of association (REF: reference allele, AF: allele frequency of alternate (ALT) allele across the pangenome). The beta value represents the effect size of a genetic variant on the expression level of the gene.

| Gene name | SNP ID | REF (G4Hunter Score) | ALT (G4Hunter Score) | TSS distance | AF | beta | q-value |
| --- | --- | --- | --- | --- | --- | --- | --- |
| *POGLUT3* | CHM13#0#chr11:108505942:>20722>20725 | G (1.56) | A (1.15) | 143 bp upstream | 0.167 | -0.726 | 9.32E-06 |
| *MEGF9* | CHM13#0#chr9:132910828:>33784>33787 | A (-) | G (1.54) | 204 bp downstream | 0.417 | -0.553 | 3.42E-05 |
| *PLCB2* | CHM13#0#chr15:38114877:>4393>4396 | C (3.43) | G (3.07) | 62 bp downstream | 0.32 | 0.424 | 1.58E-03 |
| *SAAL1* | CHM13#0#chr11:18201553:>7476>7478 | CCGG (-) | C (1.52) | 74 bp upstream | 0.864 | 0.598 | 6.86E-03 |
| *DGCR6* | CHM13#0#chr22:19281846:>2133>2135 | CGCGGGCGGG (2.06) | C (-) | 265 bp downstream | 0.044 | -0.902 | 9.23E-03 |
| *MOCS2* | CHM13#0#chr5:53937333:>41192>41194 | C (2.45) | CCGGGGG (3.35) | 210 bp upstream | 0.289 | -0.412 | 9.41E-03 |

Overall, our analyses of long-read expression data provide evidence of an association between genetic variation in pG4s proximal to TSSs and gene expression levels for several genes. The precise role of G4 formation in regulating transcription levels across the human pangenome requires further experimental validation.

### pG4 variation at or near splice site junctions are associated with alternative splicing

We next evaluated whether pG4 variation at splice-site junctions (or their vicinity) contributes to generating the transcript isoform repertoire across the human pangenome. As an initial screen, we identified a total of 39,383 pG4s near splice site junctions (pG4 overlapping within ±50 bp from the splice site, later called “splice-site pG4s”) in the human T2T reference genome (CHM13). We next studied genome-wide splice-site genetic variants with an sQTL (splicing Quantitative Trait Loci) approach (using the window size of 10 kb), utilizing the long-read PacBio transcriptome data from lymphoblastoid cell lines across the human pangenome (Human Pangenome Reference Consortium 2026; Marin and Li 2026). Among the genome-wide variants with significant correlation (*q*-value < 0.01) to splice-junction usage genome-wide (i.e., splice-site phenotypes, or the normalized proportion of retained vs. the sum of excised and retained isoforms for each exon-intron junction) identified with this approach, 1,528 were associated with 10,621 genetic variants overlapping 446 splice-site pG4s across 394 genes. To identify a more confident candidate list of genes with splicing potentially regulated by G4s, we kept only the genes where the most significant genetic variant associated with the most significant splice-site phenotype (minimum *p*-value) was within a pG4, leading to a total of 116 genes. Further, we conservatively removed any gene with a non-pG4 variant that had a higher or similar association (difference ≤ 0.5) as compared to a splice-site pG4 variant. This led to the final set of 66 candidate genes with 269 splice-site phenotypes, which were sorted by the greatest change in *p*-value between the pG4 and non-pG4 variants of that gene. Below we discuss in more detail some of the splice-site phenotypes and the associated pG4 genetic variation in genes that have at least 10^10^ times higher *p*-value at the splice-site pG4 variant as compared to the linked non-pG4 variants.

In several instances, alternative splicing was associated with pG4s located on the template strand. For example, a pG4 on the negative strand of the reference of the *ULK3* (Unc-51 Like Kinase 3) gene displayed a significant positive correlation (β: 1.620, nominal *p*-value: 3.16×10^-116^) with the intron excision (chr15:74,837,436-74,837,756) at the splice site 18 bp downstream from the center of the pG4 (Fig. 5A, C). At the same time, the reduction in stability of pG4, marked by the single nucleotide variant, C>T, was negatively associated with the excision of a slightly shorter region (chr15:74,837,436-74,837,750) 12 bps downstream, with high significance (β: -1.674, nominal *p*-value: 1.97×10^-121^; Fig. 5B, C). In other instances, alternative splicing was associated with pG4s located on the non-template strand. For example, the presence of a splice-site pG4 on the non-template strand of the *H6PD* (Hexose-6-Phosphate Dehydrogenase) gene was positively correlated (β: 1.662, nominal *p*-value: 6.84×10^-71^) with the excision of the region chr1:9,235,067-9,244,932 (Fig. 5D, F). This pG4 (the center of its annotation boundaries) is located 22 bps downstream from the splice site. The reduction in its stability due to a single nucleotide variant, C>T, was negatively correlated (β: -2.049, nominal *p*-value: 1.87×10^-54^) with the excision of a shorter region, chr1:9,235,067-9,244,924 (Fig. 5E, F). Figure S11 shows several other examples of pG4 linked variants significantly associated with splice-site phenotypes, across both strands.

**Figure 5.**
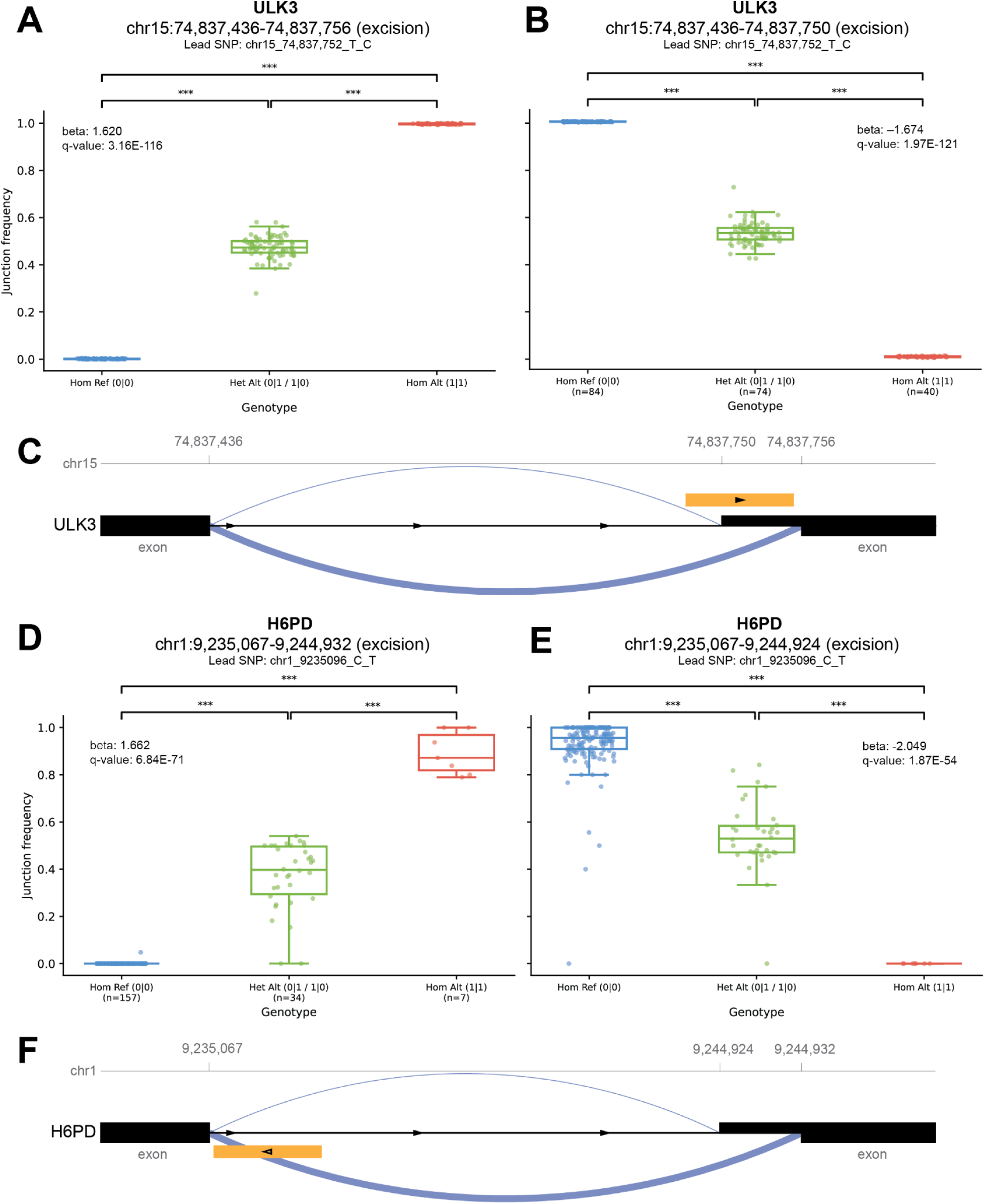
pG4-dependent splicing levels, across two candidate genes. (**A-B**) The boxplots show two different splice-site phenotypes at the *ULK3* gene associated with the variant, T>C, at chr15:74,837,752, located within the pG4. (**C**) The schematic highlights the presence of pG4 the template strand, with arrows indicating the direction of the gene. The thicker blue line represents a positive association with the pG4 bearing the C nucleotide in its positive strand, while the thinner line represents a negative association. (**D-E**) The boxplots show two different splice-site phenotypes at the ULK3 gene associated with the variant, C>T, at chr1:9,235,096 located within the pG4. (**F**) Unlike the above case, the pG4 in this case is present on the non-template strand. The thicker line represents the association with the C-bearing pG4 in its positive strand. Significance stars denote results from two-sided Mann-Whitney-Wilcoxon pairwise tests. (non-significant (ns): *p-value >* 0.05; *: 0.01 < *p-value* ≤ 0.05; **: 0.001 < *p-value* ≤ 0.01; ***: 1.00e-04 < *p-value* ≤ 0.001; ****: *p-value* ≤ 1.00e-04).

Overall, our analyses of long-read expression data indicate an association between splice-site pG4 genetic variation and isoforms for multiple genes, suggesting the G4 regulatory potential of splicing within the human pangenome. These results should be further validated in wet-lab experiments.

## DISCUSSION

### pG4s in the human pangenome

Leveraging the recent HPRC release 2 dataset (Human Pangenome Reference Consortium 2026), we predicted G4 motifs (pG4s) to study their value as population genetic markers, the selection pressure they are subject to, and their potential in gene expression regulation. In total, pG4s cover ∼19.58 Mb, or 0.65%, of the human genome (average across the human pangenome haplotypes (Human Pangenome Reference Consortium 2026). The consistency of pG4 counts per haplotype, averaging 7.58×10⁵, closely mirrored our earlier estimate of 7.69×10⁵ from the human CHM13 T2T assembly (Mohanty et al. 2025), and suggests that pG4 motif density is broadly conserved across human genomes. Notably, these values are higher than those reported in previous studies on preceding T2T assemblies (Chambers et al. 2015; Marsico et al. 2019; Tu et al. 2021).

We found that variation in pG4 presence/absence alone recapitulates the global human population structure described in previous studies (Reed and Tishkoff 2006; Campbell and Tishkoff 2010). The correspondence with canonical SNP-based analyses, including the greater diversity of African populations, the tight clustering of East Asian haplotypes, and the intermediate positioning of admixed American populations—a pattern remarkably similar to the genome-wide patterns observed in the 1000 Genomes Consortium Data (Gaspar and Breen 2019)—implies that the gain and loss of pG4-forming sequences is governed by the same demographic forces shaping broader genome evolution. This is not entirely surprising given that pG4 presence/absence is, in most cases, driven by SNVs undergoing the same mutation processes as outside of pG4s. Similarly, the fact that sequence-level variation within pG4s reproduces the genome-wide PCA structure suggests these regions are not outliers in the human diversity landscape, but are an integral part of it. What is somewhat surprising is that the population structure could indeed be recapitulated from pG4 presence/absence and pG4 sequence variation despite the elevated mutation rate at these loci (Guiblet et al. 2018; Guiblet, Cremona, et al. 2021).

Additionally, pG4s uncovered a nuanced demographic history for some human populations. In particular, the Peruvian population stood out, with the majority of its haplotypes clustering into a single clade. This observation aligns with previously documented bottleneck events in certain Latin American groups (Borda et al. 2024), which likely resulted in the observed reduction in genetic diversity.

### Selection at pG4s

Our analysis of variants within pG4s across the human genome identified a consistent pattern of selection pressure: for most types of functional regions (all but CDS), variants within pG4s are subject to stronger purifying selection than variants outside of pG4s. The most significant outcome of this research is the quantification of the difference in selection coefficients between pG4 and non-pG4 sequences across a variety of functional regions, including promoters, introns, and enhancers, and also likely UTRs. This implies that the constraint we detect is not simply a byproduct of being embedded within a functional region, but reflects selection acting on the pG4s themselves, strongly supporting their functional role. Our results are consistent with G4 function at promoters—e.g., in gene activation (Li et al. 2023), transcription factor binding (Lago et al. 2021), and activation of alternative promoters (Zhang and Mergny 2025); at introns—e.g., in alternative splicing (Georgakopoulos-Soares, Parada, Wong, et al. 2022); and at enhancers—e.g., as mediators of interactions with promoters (DeMeis et al. 2025). The selection coefficients at UTRs, albeit with low statistical confidence in several populations (Fig. S8), suggest how pG4s may contribute at 5’ UTRs, e.g., as translational enhancers (Lee et al. 2024) or repressors (Beaudoin and Perreault 2010), and at 3’ UTRs, e.g., in alternative polyadenylation and mRNA shortening (Beaudoin and Perreault 2013; D. S. M. Lee et al. 2020; Rouleau et al. 2017).

The strong selection on pG4s at replication origins—contrasting sharply with the near-neutral non-G4 flanking sequences—provides population-genetic support for the functional importance of pG4s located at Origin G-rich Repeated Elements (OGREs) in replication initiation, a mechanism previously inferred from molecular data alone (Besnard et al. 2012; Langley et al. 2016).

Our data confirm strong selection acting on protein-coding regions as a whole. However, in contrast with other types of functional regions, selection pressure is stronger at non-pG4 sequences than at pG4 sequences within protein-coding regions. This might indicate selection against pG4 formation specifically in these regions, consistent with our prior results (Guiblet, DeGiorgio, et al. 2021). It was suggested that such selection occurs through the use of particular codons (Mirihana Arachchilage et al. 2019). Subsetting the coding regions to 0-fold and 4-fold sites yielded even starker results. Selection pressure on the former, which constitute purely nonsynonymous sites, was even stronger than on coding regions as a whole, i.e., the highest among all regions tested. Although we could not quantify selection pressure at 4-fold sites with statistical confidence, these sites displayed near-neutral coefficients both at pG4s and outside them, as expected due to codon degeneracy at these sites, resulting from synonymous mutations.

The broad concordance of selection coefficients at pG4s, despite the demographic and geographic diversity of the KHV, PEL, JPT, and GWD populations, suggests that evolutionary pressures acting on these motifs may have originated prior to recent human lineage divergence. This suggests that pG4s critical for function are evolutionarily old—with higher sharing and stronger selection signatures, consistent with results in another recent study by our group (Zhang et al. 2026).

Notwithstanding this plethora of evidence, our analyses have several limitations that warrant attention. First, our inferences rely on computationally predicted G4 motifs; a substantial fraction of these may not form stable G4s *in vivo* under physiological conditions (Kouzine et al. 2019; Sieg et al. 2025). Our selection estimates are thus, in effect, averaged over sequences that may or may not form G4s. This means we are likely underestimating the true constraint on *bona fide* G4-forming sites. Second, the low statistical confidence in our inferences across 4-fold degenerate sites and UTR regions—stemming from the limited number sites within pG4s—means that the corresponding estimates must be interpreted with caution. Third, the choice of NGNR regions as a neutral reference, while standard (Guiblet, Cremona, et al. 2021; Arbiza et al. 2012), assumes these regions evolve under no selective constraint, which may not always be true.

### pG4s as regulators of alternative splicing and gene expression

Our findings also highlight an association between pG4 variation and gene expression across a diverse human pangenome cohort, reinforcing the emerging consensus that G4 structures serve as active cis-regulatory elements (Robinson et al. 2021; Du et al. 2008; Rhodes and Lipps 2015). Using long-read sequencing data and haplotype-resolved expression profiling, we found that pG4 variation at gene promoters is significantly associated with transcriptional output in a gene-specific manner, with some pG4s acting as transcriptional activators and others as repressors. This result supports prior observations of pG4s playing both enhancing (C.-Y. Lee et al. 2020) and repressive (Broxson et al. 2011; Smestad and Maher 2015) roles in regulating transcription levels. SNPs in cis-regulatory elements, such as promoters, were previously shown to lead to intermediate phenotypes in heterozygotes (i.e., codominance) (Jones and Swallow 2011). The observation of intermediate expression levels in individuals heterozygous for the presence/absence of pG4 further supports codominance at cis-regulatory loci, suggesting that pG4-mediated regulation operates in a dosage-sensitive fashion.

We found a significant association between pG4 motifs at splice sites and alternative splicing of genes. This is consistent with previous studies showing enrichment of pG4s at intron-exon junctions (Georgakopoulos-Soares, Parada, Wong, et al. 2022) and supporting their putative role in splicing through different mechanisms (Peynet et al. 2026; Varshney et al. 2020). By utilizing pangenome sequences and long-read expression data, as well as sQTL analysis, we were, for the first time, able to pinpoint pG4s regulating alternative splicing in a locus-specific manner in human populations. Notably, we also found pG4s located on either strand associated significantly with the splice-site phenotypes, suggesting both-strand pG4s might be required for alternative splicing, although the underlying mechanism of regulation might be different as described above.

Although promising, we note that, first, our analysis is limited to lymphoblastoid cell lines and that G4 effects on gene expression and splicing may differ across tissue types and developmental stages. Second, while the QTL framework with correction for multiple hits reduces the risk of false discovery, larger cohorts would improve statistical power to detect pG4-expression and pG4-isoform associations with modest effect sizes, thereby increasing the candidate gene set. Third, the eQTL analysis is sensitive to RNA sequencing depth, which needs to be considered (Human Pangenome Reference Consortium 2026). Higher sequencing depth is expected to uncover additional significant associations. Fourth, our QTL results have passed through multiple filtering steps, and thus should be considered conservative. Finally, *in vitro* and *in vivo* experiments will be needed to confirm G4 formation and disruption, and their influence on gene expression and alternative splicing, as pG4 scoring methods only predict G4 formation.

## METHODS

### Prediction and inference of the presence/absence variation for pG4s in the human pangenome

The FASTA files for 234 genomes comprising 466 haplotypes were downloaded from the release2 v1.0 of the HPRC consortium (https://github.com/human-pangenomics/hprc_intermediate_assembly/blob/ee0fd4cb216e4bdc0d13d17510ea55548c7f4c0f/data_tables/assemblies_release2_v1.0.index.csv). Using the G4Discovery (Mohanty et al. 2025) tool compatible with the PanSN naming convention (https://github.com/pangenome/PanSN-spec), G4s were predicted for each of the 466 haplotypes. The `G4Discovery` tool combines pqsfinder (Hon et al. 2017) and G4Hunter (Bedrat et al. 2016) scoring. Candidate pG4s are then filtered by minimum tetrad count and score thresholds for both tools, and deduplicated by selecting the highest-scoring, shortest G4 per genomic position, producing a non-overlapping BED file output for each strand.

To derive reference-independent data on pG4s presence, the analysis was performed separately for each chromosome of each haplotype. To account for different naming conventions across assemblies, scaffolds were labeled with chromosome numbers using the file at https://github.com/human-pangenomics/hprc_intermediate_assembly/blob/main/data_tables/annotation/chrom_assignment/chrom_alias_hprc_r2_v1.0.index.csv. Some of the contigs and scaffolds in some haplotypes lacked chromosome assignment and were hence labeled as “unassigned.” Homologous pG4 regions were identified across all 466 haplotypes by using pG4 annotations from a selected set of haplotypes (CHM13 and five additional paternal haplotypes from HG002, HG00597, HG01358, HG02572, and HG04184), which represent diverse superpopulations. This derivation was performed on a per-chromosome basis using implicit pangenome (`IMPG:` https://github.com/pangenome/impg) in the pairwise alignment file (`25,221.PAF`) generated from the HPRCv2 genomes (https://github.com/pangenome/HPRCv2) via `WFMASH` (Guarracino et al. 2026). The pG4s from other haplotypes that were not included in the homologous region catalog were then extracted. `IMPG` was independently rerun on the newly extracted set, using the same `PAF` file. The resulting data were subsequently combined with the previously derived set from the selected haplotypes. We converted the `BEDPE` file to `BED` format using a custom Python script (https://github.com/makovalab-psu/panG4nome) built on the `igraph` module (Csardi and Nepusz 2006). In this approach, each pair of homologous regions was treated as an edge connecting two nodes, and the connected components of the resulting graph were identified. Each node (i.e., each homologous region) was then assigned a unique region name. For each haplotype, we determined the exact location of pG4s surrounding the labeled regions by overlapping these regions with a chromosome-level BED file containing pG4s from all haplotypes.

To ensure unbiased and error-free downstream analysis, pG4s located in unassigned scaffold regions were iteratively removed until convergence, ensuring no residual overlap with excluded regions. In addition, to achieve a high-confidence dataset, we removed flagged regions reported by two independent flagging methods. The first method, based on `NucFlag` (https://github.com/logsdon-lab/NucFlag), was applied separately to ONT and HiFi reads, and the second method utilized `HMMFlagger` (Asri et al. 2026). For our analysis, a region was designated as "flagged" only if it was flagged in at least two of these three datasets—`NucFlag` ONT, `NucFlag` HiFi, and `HMMFlagger`—in more than 1% of the haplotypes, in order to retain only the well-resolved regions of the assemblies. The removal was performed through a second iterative filtering step.

The process resulted in a high-confidence pG4 dataset, which was then converted into a presence/absence matrix for all haplotypes. To construct this pG4 presence/absence matrix across haplotypes for each chromosome, pG4s overlapping multiple pangenome graph regions were first linked into a graph of connected components. These connected regions were then consolidated under a single representative label. Unique identifiers were subsequently assigned to pG4s absent from the pangenome alignment. The final output included a reference `BED` file containing non-redundant pG4 loci and a presence/absence matrix, with a collection of pG4s at orthologous locations across human haplotypes termed as a “pG4 family”. The output presence/absence matrix quantified the number of pG4s per pG4 family. A schematic of this method is shown in Fig. S12.

### Maximum parsimony phylogenetic tree construction from pG4 presence/absence

Maximum parsimony phylogenetic analysis of the pG4 presence/absence matrix was performed using PAUP* (Swofford 2003) across the haplotypes in the human pangenome. Prior to the analysis, the data were binarized because the non-binarized matrix contained the number of pG4s per pG4 family per sample, which could exceed 1 due to duplicated regions. Following conversion of the binarized PAV matrix to a `nexus`-formatted alignment, a heuristic search was conducted with 10 replicates using the "as-is" sequence addition order, retaining up to 100 trees in memory. The first most parsimonious tree recovered by the heuristic search was saved in Newick format. The unrooted tree was constructed using the `ggtree` (Yu 2020) package in R.

### Quantifying variation at pG4 and non-pG4 regions

To characterize nucleotide variation within and around pG4 regions, we extracted all pG4 regions in the presence/absence matrix and discarded those lacking orthologous intervals in the CHM13 genome. Following this, regions containing a CHM13 pG4 were extended to include the corresponding pG4 intervals. The other regions, devoid of CHM13 pG4s, were retained.

Together, these regions were defined as “pG4” regions. SNPs overlapping the extended regions were extracted from a whole-genome `PGGB`-graph variant file (20250603_hprc8424.CHM13.laced.norm.vcf.gz, downloaded from https://github.com/pangenome/HPRCv2), using `bcftools` (Danecek et al. 2021) and retaining only biallelic SNVs.

For functional annotations, genome annotation files in General Feature Format (`GFF`) for CHM13 were downloaded from `NCBI` (O’Leary et al. 2024), similar to (Mohanty et al. 2025). The longest transcripts for protein-coding genes were extracted using the script provided on GitHub (https://gist.github.com/karolpal-jr/48213a0a65475e44f708d5d815127bc3). From these, introns were identified as non-exonic mRNA segments, and UTRs (5’ and 3’) were identified as the non-coding portions of exons. Promoters were standardized as 1-kb regions upstream of transcription start sites. Regions without any overlap with genes, promoters, origins of replication, enhancers, and repeats were used to form the NGNR “neutral” benchmark. Core and stochastic origin annotations were taken from (Akerman et al. 2020) and transformed to CHM13 coordinates using the UCSC `liftOver` tool (Hinrichs et al. 2006). A comprehensive `VCF` was generated for all of these genomic compartments—including promoters, UTRs, CDS, introns, enhancers, replication origins, and NGNR regions—and all pG4 intervals were removed from it. Together, these regions were defined as “non-pG4” regions. The CDS were subsetted as 4-fold and 0-fold degenerate sites using the `Degeneracy` tool (https://github.com/zhangrengang/degeneracy).

### Ancestral inference using chimpanzee and orangutan genomes as outgroups

For ancestral inference, we used two outgroup species—chimpanzee (*Pan troglodytes*) and Bornean orangutan (*Pongo pygmaeus*). The pairwise alignments between the complete genomes of each of these species and the human CHM13 (Mohanty et al. 2025) were used to extract the corresponding outgroup nucleotide for each biallelic SNV, using the `bx.align` package (https://bx-python.readthedocs.io/en/latest/lib/bx.align.html). Ancestral sites were inferred using a method that requires both outgroup nucleotides to match either the reference or alternate allele (https://github.com/makovalab-psu/panG4nome); otherwise, the ancestral state was not inferred. The inferred ancestry was later added as the `INFO/AAS` tag into the `VCF`s using `bcftools annotate` (Danecek et al. 2021).

### Principal Component Analysis using the variation data

To assess population structure across haplotypes, we performed PCA on the pG4 variation data. The variant call set was first filtered to retain only biallelic SNPs using `bcftools view` (Danecek et al. 2021). The resulting VCF was imported into `PLINK2` (Chang et al. 2015), where genotypes with only one allele successfully called (half-calls) were encoded as missing. Prior to PCA, variants were pruned for linkage disequilibrium (LD) using a sliding window of 50 variants, a step size of 5, and an r^2^ threshold of 0.1. The LD-pruned variant set was then used to compute the top 20 principal components in `PLINK2`. Finally, the PC1 vs PC2 plot was marked with population and superpopulation identifiers.

### Quantifying selection coefficients inside and outside of pG4s at various functional regions using fastDFE

To facilitate robust DFE analysis, we selected four distinct populations—GWD, JPT, PEL, and KHV (see Results for a justification of this choice). To determine the anticipated site counts for a new sample size of individuals, the subsampling process employs a hypergeometric projection in "probabilistic" mode, based on the original counts. This also smoothens the bins to fit the model better. Our workflow for each population began by reducing the pG4 and non-pG4 (as defined previously) `VCF` files to include only the individual samples belonging to that population. We then extracted dimorphic alleles labeled with ancestral states and excluded all monomorphic regions. Since DFE inference requires counts of monomorphic sites, these were recovered from the original `PGGB` graph-based `VCF`. Finally, to construct the site-frequency spectra (SFS), we partitioned the pG4 and non-pG4 BED files into specific functional genomic regions as defined previously.

The underlying SFS were unfolded using ancestral labels to identify derived allele frequencies, with monomorphic site counts incorporated at the beginning of the distribution. Sex chromosomes were excluded from the estimation process due to their different ploidy levels. To determine the population-scaled deleterious selection coefficient S_d_ (*S_d_ = 4N_e_s*), we utilized the *BaseInference* module of `fastDFE` (Sendrowski and Bataillon 2024) employing

Gamma-Exponential Parametrization, subsampling to 25 haplotypes, and setting the number of runs to 30. Since, to estimate the population demography, `fastDFE` requires selected and neutral SFS for “pG4” regions, the NGNR pG4 SFS was used as the neutral baseline, and similarly, for “non-pG4” regions, the NGNR non-pG4 SFS was used as the neutral baseline. This was done to control for differences in mutation rates between pG4 and non-pG4 sites. In this analysis, as described in the `fastDFE` manual, the probability of beneficial selection was set to zero (*p_b_ = 0*), the beneficial population-scaled selection coefficient to one (*Sb = 1*), and the dominance coefficient to 0.5 (*h = 0.5*), assuming semi-dominant mutations. The gamma parametrization integrates gamma and exponential components to model deleterious and beneficial coefficients. However, by setting the probability of beneficial selection to zero, we removed the exponential component. Therefore, using the gamma-parameterized model, we estimated the deleterious population-scaled selection coefficient (*S_d_*), the shape factor for the gamma distribution, and the ancestral-allele misidentification rate (*eps*).

To assess the stability of each *S_d_* estimate, we performed 30 independent estimation runs. Estimates for a run were considered unstable if a run yielded a log-likelihood value more than 1 unit lower than the best run. They were excluded from confidence interval calculations. Model fit to the observed SFS was evaluated using the normalized L1 residual from fastDFE; values exceeding 0.1 or with a standard deviation ≥15 for the *S_d_* estimate were classified as a poor fit and considered low confidence. Finally, absolute deleterious selection coefficients were categorized into three distinct regimes based on the following boundary definitions (Johri et al. 2020): neutral selection, *S_d_ < 1*; weakly deleterious selection, *1 ≤ S_d_ < 10*; and moderately/strongly deleterious selection, *S_d_ ≥ 10*.

### eQTL analysis

Kinnex (PacBio long-read RNA-seq) data were produced for 206 HPRC samples from the lymphoblast cell line of each donor (Marin and Li 2026). Full-length non-chimeric (FLNC) reads produced by the PacBio `Iso-Seq` pipeline (https://github.com/PacificBiosciences/IsoSeq) were clustered into representative consensus sequences using the `Iso-Seq` cluster tool (https://github.com/PacificBiosciences/IsoSeq), retaining only clusters supported by two or more FLNC reads (i.e., singletons were excluded). Representative transcript cluster sequences were then processed with `IsoQuant` (Prjibelski et al. 2023) against the GRCh38 reference genome and the GENCODE v44 comprehensive annotation, which assigned each cluster to an annotated gene identifier. Gene-level expression level quantification was obtained by summing the FLNC read counts of all clusters assigned to a given gene. To enable cross-sample comparisons, gene-level counts per sample were normalized to counts per million (CPM).

Genome annotation files in `GFF` for CHM13 were downloaded from NCBI (O’Leary et al. 2024). From these annotations, we extracted the longest corresponding transcript, CDS, and exon entries for protein-coding genes (using the snippet available at https://github.com/makovalab-psu/panG4nome). For each gene, we then identified the TSS positions of the longest transcript reported for the CHM13 reference assembly. To account for population structure or LD in phenotypes, we calculate covariates by extracting all the variants in this region with `bcftools view` (Danecek et al. 2021) and retaining the 206 samples present in the Kinnex dataset. Using PLINK2 (Chang et al. 2015), we computed 20 principal component eigenvectors, with genotypes with only one successfully called allele (half-calls) encoded as missing. Using the CPM values for protein-coding genes in our dataset, we filtered out genes with low expression levels by setting a minimum threshold of 1 CPM in at least 10 samples. This procedure removed 7,423 genes out of 19,187 genes. A nominal eQTL analysis was performed using tensorQTL (Taylor-Weiner et al. 2019), with a window size of 20 kb, following the inverse-normal transformation of the CPM gene expression data. Sex chromosomes were removed from this analysis.

Significant variants were filtered to include only those within 3 kb (2 kb upstream to 1 kb downstream) of the TSS of the longest transcript of each protein-coding gene. These variants were further intersected with pG4s and pG4 orthologs in the CHM13 genomic coordinates using bedtools intersect (Quinlan and Hall 2010). The reference and alternate alleles for these pG4 variants were inferred from the VCF, and their predicted effects on G4 stability and formation were assessed. For the pairwise statistical comparison between pG4 zygosity and log-normalized gene expression levels (Fig. 4), the statannotations (Charlier et al. 2022)

Python module was used, with the two-sided Mann-Whitney-Wilcoxon test with the Benjamini-Hochberg correction.

### sQTL analysis

Splice junction counts were extracted from the FLNC read alignments using *`scmatrix`* (Huang and Li 2026), which clusters and quantifies intron excision events in a manner analogous to `LeafCutter` (Li et al. 2018). Junction cluster usage ratios were computed across all samples and quantile-quantile (QQ) normalized to generate a per-cluster phenotype matrix. sQTL analysis was performed using `rust-fastQTL` (https://github.com/huangnengCSU/rust-fastqtl), a Rust implementation of the `FastQTL` permutation framework. The variant input consisted of a merged VCF combining biallelic SNPs from the HPRC Minigraph-Cactus `DipCall` calls with per-sample pG4 presence/absence encoded as binary variants (Human Pangenome Reference Consortium 2026), allowing joint testing of both conventional genetic variation and pG4 occupancy against splicing phenotypes. Only variants with a minor allele frequency ≥0.05 and present in at least 10 samples were considered. Association testing was performed within a 10-kb cis-window around each junction cluster, using a permutation pass (1,000–10,000 permutations) to calibrate per-phenotype null distributions, followed by a nominal pass to recover all tested variant–junction pairs. The first 10 principal components of the QQ-normalized phenotype matrix were included as covariates to account for global splicing variation across samples.

Genome-wide FDR correction was applied to permutation-pass beta-approximated *p*-values using the standard `FastQTL` two-step procedure, and sQTL associations were considered significant at FDR < 0.01. Splice sites were obtained from genes annotated in CHM13, conservatively limiting to genes with a known orthology to other great ape species in NCBI (using the flag ---ortholog 9604 in NCBI’s software Datasets (v18.9.1) (O’Leary et al. 2024). Since the exact splice junctions were not provided in the annotation, we extracted 4 bp around every exon (2 bp from the start/end of the exon and 2 bp from the adjacent intron, leaving out the start of the first exon and end of the last exon for every transcript). Subsequently, we identified pG4s located within 50 bp upstream and downstream of splice sites, known as “splice-site pG4s”. We then mapped these pG4s to GRCh38 coordinates using the UCSC `liftOver` tool. Next, we overlapped all significant variants from the sQTL analysis with the pG4s using `bedtools intersect`. We identified all significant sQTLs or splice-site phenotypes correlated with splice-site pG4s. To ensure that the pG4 variant was the lead variant (i.e., had the lowest *p*-value) in the most significant sQTL for each gene, we removed all genes that did not meet this condition. The `statannotations` Python module was utilized for pairwise statistical evaluations between pG4 zygosity and log-normalized gene expression (Fig. 4), employing a two-sided Mann-Whitney-Wilcoxon test and applying Benjamini-Hochberg correction.

## DECLARATIONS

### DATA AND CODE AVAILABILITY

All source code is available on GitHub at https://github.com/makovalab-psu/panG4nome.

## ACKNOWLEDGMENTS

We are grateful to Neng Huang, Heng Li, Karen Miga, Karol Pal, and Jacob Sieg for discussing the results and providing useful suggestions. The authors also recognize the Penn State Institute for Computational and Data Sciences (ICDS) (RRID:SCR_025154) for providing access to computational research infrastructure (RRID:SCR_026424).

## Human Pangenome Reference Consortium Version 2 Authors

Derek Albracht^1^, Ivan A. Alexandrov^2^, Jamie Allen^3^, Alawi A. Alsheikh-Ali^4^, Nicolas Altemose^5^, Casey Andrews^6^, Dmitry Antipov^7^, Lucinda Antonacci-Fulton^1^, Mobin Asri^8^, Marcelo Ayllon^9^, Jennifer R. Balacco^10^, Floris P. Barthel^11^, Edward A. Belter Jr^1^, Halle D. Bender^8^, Andrew P. Blair^8^, Davide Bolognini^12^, Katherine E. Bonini^13^, Christina Boucher^14^, Guillaume Bourque^15,16,17^, Silvia Buonaiuto^18^, Shuo Cao^18^, Andrew Carroll^19^, Ann M. Mc Cartney^8^, Monika Cechova^8^, Mark J.P. Chaisson^20^, Pi-Chuan Chang^19^, Xian Chang^8^, Jitender Cheema^3^, Haoyu Cheng^21^, Claudio Ciofi^22^, Hiram Clawson^8^, Sarah Cody^1^, Vincenza Colonna^18^, Holland C. Conwell^23^, Robert Cook-Deegan^24^, Mark Diekhans^8^, Maria Angela Diroma^22^, Daniel Doerr^25,26,27^, Zheng Dong^6^, Danilo Dubocanin^5^, Richard Durbin^28,29^, Jana Ebler^25,30^, Evan E. Eichler^9,31^, Jordan M. Eizenga^8^, Parsa Eskandar^8^, Eddie Ferro^14^, Anna-Sophie Fiston-Lavier^32,33^, Sarah M. Ford^23^, Willard W. Ford^34^, Giulio Formenti^10^, Adam Frankish^3^, Mallory A. Freeberg^3^, Qichen Fu^6^, Stephanie M. Fullerton^35^, Robert S. Fulton^1^, Yan Gao^36^, Gage H. Garcia^9^, Obed A. Garcia^37^, Joshua M.V. Gardner^8^, Shilpa Garg^38^, Erik Garrison^18^, Nanibaa’ A. Garrison^39,40,41^, John E. Garza^1^, Margarita Geleta^42^, Mohammadmersad Ghorbani^43^, Tina A. Graves-Lindsay^1^, Richard E. Green^23^, Cristian Groza^44^, Bida Gu^20^, Andrea Guarracino^11,18^, Melissa Gymrek^45^, Maximilian Haeussler^8^, Leanne Haggerty^3^, Ira M. Hall^46,47^, Nancy F. Hansen^7^, Yue Hao^11^, Mohammad Amiruddin Hashmi^4^, David Haussler^8^, Prajna Hebbar^8^, Peter Heringer^25,26,27^, Glenn Hickey^8^, Todd L. Hillaker^8^, S. Nakib Hossain^3^, Neng Huang^36,48^, Sarah E. Hunt^3^, Toby Hunt^3^, Alexander G. Ioannidis^5,8^, Nafiseh Jafarzadeh^8^, Nivesh Jain^10^, Erich D. Jarvis^10,31^, Maryam Jehangir^11^, Juan Jiang^6^, Eimear E. Kenny^13^, Juhyun Kim^7^, Bonhwang Koo^10^, Sergey Koren^7^, Milinn Kremitzki^1,6^, Charles H. Langley^49^, Ben Langmead^50^, Heather A. Lawson^6^, Daofeng Li^6^, Heng Li^36,48^, Wen-Wei Liao^46,47^, Jiadong Lin^9^, Tianjie Liu^6^, Glennis A. Logsdon^51^, Ryan Lorig-Roach^8^, Jonathan LoTempio Jr^52^, Hailey Loucks^8^, Jane E. Loveland^3^, Jianguo Lu^53^, Shuangjia Lu^46,47^, Julian K. Lucas^8^, Walfred Ma^20^, Juan F. Macias-Velasco^1,6,54^, Kateryna D. Makova^55^, Maximillian G. Marin^36,48^, Christopher Markovic^1^, Tobias Marschall^25,30^, Franco L. Marsico^18^, Fergal J. Martin^3^, Mira Mastoras^8^, Capucine Mayoud^32^, Brandy McNulty^8^, Jack A. Medico^10^, Julian M. Menendez^8^, Karen H. Miga^8^, Anna Minkina^56^, Matthew W. Mitchell^57^, Saswat K. Mohanty^58^, Younes Mokrab^43,59,60^, Jean Monlong^61^, Shabir Moosa^43^, Avelina Moreno-Ochando^62,63^, Shinichi Morishita^64^, Jonathan M. Mudge^3^, Katherine M. Munson^9^, Njagi Mwaniki^65^, Nasna Nassir^4^, Chiara Natali^22^, Shloka Negi^8^, Lingbin Ni^9^, Adam M. Novak^8^, Faith Okamoto^8^, Pilar N. Ossorio^66^, Chie Owa^64^, Sadye Paez^10^, Benedict Paten^8^, Clelia Peano^12,67^, Adam M. Phillippy^7^, Brandon D. Pickett^7^, Laura Pignata^18^, Nadia Pisanti^65^, David Porubsky^9,68^, Pjotr Prins^18^, Anandi Radhakrishnan^8^, T. Rhyker Ranallo-Benavidez^11^, Brian J. Raney^8^, Mikko Rautiainen^69^, Alessandro Raveane^12^, Luyao Ren^9,31^, Arang Rhie^7^, Fedor Ryabov^70,71^, Samuel Sacco^23^, Farnaz Salehi^18^, Michael C. Schatz^50,72^, Laura B. Scheinfeldt^73^, Aarushi Sehgal^34^, William E. Seligmann^23^, Mahsa Shabani^74^, Kishwar Shafin^19^, Shadi Shahatit^32^, Ruhollah Shemirani^13^, Vikram S. Shivakumar^50^, Swati Sinha^3^, Jouni Sirén^8^, Linnéa Smeds^58^, Steven J. Solar^7^, Marco Sollitto^10,22^, Nicole Soranzo^12,28,75^, Andrew B. Stergachis^9,56^, Marie-Marthe Suner^3^, Yoshihiko Suzuki^64^, Arda Söylev^25,30^, Ahmad Abou Tayoun^76,77^, Jack A.S. Tierney^3^, Chad Tomlinson^1^, Francesca Floriana Tricomi^3^, Mohammed Uddin^4,78^, Matteo Tommaso Ungaro^23,79^, Rahul Varki^14^, Flavia Villani^18^, Ivo Violich^8^, Mitchell R. Vollger,^56^, Brian P. Walenz^7^, Charles Wang^80^, Lisa E. Wang^13^, Ting Wang^1,6,54^, Aaron M. Wenger^81^, Conor V. Whelan^10^, Zilan Xin^6^, Zheng Xu^6^, Kai Ye^82^, DongAhn Yoo^9^, Wenjin Zhang^6^, Ying Zhou^36^, Xiaoyu Zhuo^6^, Giulia Zunino^12^

^1^ McDonnell Genome Institute, Washington University School of Medicine, St. Louis, MO 63108, USA

^2^ Department of Human Molecular Genetics and Biochemistry, Faculty of Medical and Health Sciences, Tel Aviv University, Tel Aviv 69978, Israel

^3^ European Molecular Biology Laboratory, European Bioinformatics Institute (EMBL-EBI), Wellcome Genome Campus, Hinxton, Cambridge CB10 1SD, UK

^4^ Center for Applied and Translational Genomics (CATG), Mohammed Bin Rashid University of Medicine and Health Sciences, Dubai Health, Dubai, UAE

^5^ Department of Genetics, Stanford University, Palo Alto, CA 94304 USA

^6^ Department of Genetics, Washington University School of Medicine, St. Louis, MO 63110, USA

^7^ Genome Informatics Section, Center for Genomics and Data Science Research, National Human Genome Research Institute, National Institutes of Health, Bethesda, MD 20892, USA

^8^ UC Santa Cruz Genomics Institute, University of California, Santa Cruz, CA 95060, USA

^9^ Department of Genome Sciences, University of Washington School of Medicine, Seattle, WA 98195, USA

^10^ The Vertebrate Genome Laboratory, The Rockefeller University, New York, NY 10065, USA

^11^ Bioinnovation and Genome Sciences, The Translational Genomics Research Institute (TGen), Phoenix, AZ 85004, USA

^12^ Human Technopole, Milan, Italy

^13^ Institute for Genomic Health, Icahn School of Medicine at Mount Sinai, New York, NY 10029, USA

^14^ Department of Computer and Information Science and Engineering, University of Florida, Gainesville, FL 32611, USA

^15^ Canadian Center for Computational Genomics, McGill University, Montréal, QC H3A 0G1, Canada

^16^ Department of Human Genetics, McGill University, Montréal, QC H3A 0G1, Canada

^17^ Victor Phillip Dahdaleh Institute of Genomic Medicine, Montréal, QC H3A 0G1, Canada

^18^ Department of Genetics, Genomics and Informatics, University of Tennessee Health Science Center, Memphis, TN 38163, USA

^19^ Google LLC, Mountain View, CA 94043, USA

^20^ Quantitative and Computational Biology, University of Southern California, Los Angeles, CA 90089, USA

^21^ Department of Biomedical Informatics and Data Science, Yale School of Medicine, New Haven, CT 06510, USA

^22^ Department of Biology, University of Florence, Sesto Fiorentino, FI 50019, Italy

^23^ Department of Ecology and Evolutionary Biology, University of California, Santa Cruz, CA 95060, USA

^24^ Arizona State University, Consortium for Science, Policy & Outcomes, Washington, DC 20006, USA

^25^ Center for Digital Medicine, Heinrich Heine University Düsseldorf, Düsseldorf, NRW, DE

^26^ Department for Endocrinology and Diabetology at the Medical Faculty and University Hospital Düsseldorf, Heinrich Heine University Düsseldorf, Düsseldorf, NRW, DE

^27^ Paul-Langerhans-Group Computational Diabetology, German Diabetes Center (DDZ) and Leibniz Institute for Diabetes Research, Düsseldorf, NRW, DE

^28^ Wellcome Sanger Institute, Genome Campus, Hinxton, CB10 1RQ, UK

^29^ Department of Genetics, University of Cambridge, Cambridge, CB2 3EH, UK

^30^ Institute for Medical Biometry and Bioinformatics, Medical Faculty and University Hospital Düsseldorf, Heinrich Heine University, Düsseldorf, NRW, DE

^31^ Howard Hughes Medical Institute, Chevy Chase, MD 20815, USA

^32^ ISEM, Univ Montpellier, CNRS, IRD, Montpellier, FR

^33^ Institut Universitaire de France, Paris, FR

^34^ Department of Computer Science and Engineering, University of California San Diego, La Jolla, CA 92093, USA

^35^ Department of Bioethics & Humanities, University of Washington School of Medicine, Seattle, WA 98195, USA

^36^ Department of Data Science, Dana-Farber Cancer Institute, Boston, MA 02215, USA

^37^ Department of Anthropology, University of Kansas, Lawrence, KS 66045, USA

^38^ School of Health Sciences, University of Manchester, Manchester M13 9PL, UK

^39^ Traditional, ancestral and unceded territory of the Gabrielino/Tongva peoples, Institute for Society & Genetics, University of California, Los Angeles, Los Angeles, CA 90095, USA

^40^ Traditional, ancestral and unceded territory of the Gabrielino/Tongva peoples, Institute for Precision Health, David Geffen School of Medicine, University of California, Los Angeles, Los Angeles, CA 90095, USA

^41^ Traditional, ancestral and unceded territory of the Gabrielino/Tongva peoples, Division of General Internal Medicine & Health Services Research, David Geffen School of Medicine, University of California, Los Angeles, Los Angeles, CA 90095, USA

^42^ Department of Electrical Engineering and Computer Science, University of California, Berkeley, Berkeley, CA 94720, USA

^43^ Medical and Population Genomics Lab, Sidra Medicine, Doha, Qatar

^44^ Montreal Heart Institute, Montréal, QC, Canada

^45^ Department of Pediatrics, University of California San Diego, La Jolla, CA 92093, USA

^46^ Center for Genomic Health, Yale University School of Medicine, New Haven, CT 06510, USA

^47^ Department of Genetics, Yale University School of Medicine, New Haven, CT 06510, USA

^48^ Department of Biomedical Informatics, Harvard Medical School, Boston, MA 02115, USA

^49^ Department of Evolution and Ecology and the Center for Population Biology, University of California, One Shields, Davis, CA 95616, USA

^50^ Department of Computer Science, Johns Hopkins University, Baltimore, MD 21218, USA

^51^ Department of Genetics, Epigenetics Institute, Perelman School of Medicine, University of Pennsylvania, Philadelphia, PA 19104, USA

^52^ Department of Pediatrics, Division of Genetics, School of Medicine, University of California, Irvine, CA 92697, USA

^53^ Sun Yat-sen University, Guangzhou, China

^54^ Edison Family Center for Genome Sciences & Systems Biology, Washington University School of Medicine, St. Louis, MO 63110, USA

^55^ Department of Biology and Center for Medical Genomics, Penn State University, University Park, PA 16802, USA

^56^ Division of Medical Genetics, Department of Medicine, University of Washington School of Medicine, Seattle, WA 98195, USA

^57^ The Jackson Laboratory for Genomic Medicine, Farmington, CT 06032, USA

^58^ Department of Biology, Penn State University, University Park, PA 16802, USA

^59^ Department of Biomedical Science, College of Health Sciences, Qatar University, Doha, Qatar

^60^ Department of Genetic Medicine, Weill Cornell Medicine-Qatar, Doha, Qatar

^61^ IRSD - Digestive Health Research Institute, University of Toulouse, INSERM, INRAE, ENVT, UPS, Toulouse, FR

^62^ MATCH biosystems, S.L., Elche, Spain

^63^ Universidad Miguel Hernández de Elche, Elche, Spain

^64^ Department of Computational Biology and Medical Sciences, The University of Tokyo, Kashiwa, Chiba 277-8561, Japan

^65^ Department of Computer Science, University of Pisa, Pisa, Italy

^66^ Law School, University of Wisconsin-Madison, Madison, WI 53706, USA

^67^ Institute of Genetics and Biomedical Research, UoS of Milan, National Research Council, Milan, Italy

^68^ Genome Biology Unit, European Molecular Biology Laboratory (EMBL), Heidelberg, DE

^69^ Institute for Molecular Medicine Finland, Helsinki Institute of Life Science, University of Helsinki, Helsinki, Finland

^70^ The Center for Bio- and Medical Technologies, Moscow, RUS

^71^ Centre for Biomedical Research and Technology, HSE University, Moscow, RUS

^72^ Department of Biology, Johns Hopkins University, Baltimore, MD 21218, USA

^73^ Coriell Institute for Medical Research, Camden, NJ 08103, USA

^74^ University of Amsterdam, Amsterdam, Netherlands

^75^ School of Clinical Medicine, University of Cambridge, Cambridge, CB2 0SP, UK

^76^ Center for Genomic Discovery, Mohammed Bin Rashid University, Dubai Health, UAE

^77^ Dubai Health Genomic Medicine Center, Dubai Health, UAE

^78^ GenomeArc Inc, Mississauga, ON, Canada

^79^ Department of Biology and Biotechnologies “Charles Darwin”, University of Rome “La Sapienza”, Rome 00185, IT

^80^ Center for Genomics, Loma Linda University School of Medicine, Loma Linda, CA 92350, USA

^81^ PacBio, Menlo Park, CA 94025, USA

^82^ The first affiliated hospital of Xi’an Jiaotong University, Xi’an Jiaotong University, Xi’an, Shaanxi, 710049, China

