## Supplementary Figures for "Variation and selection at predicted G-quadruplexes across the human pangenome"

**Supplementary Figure 1**. The average number of haplotype-specific pG4s per individual.

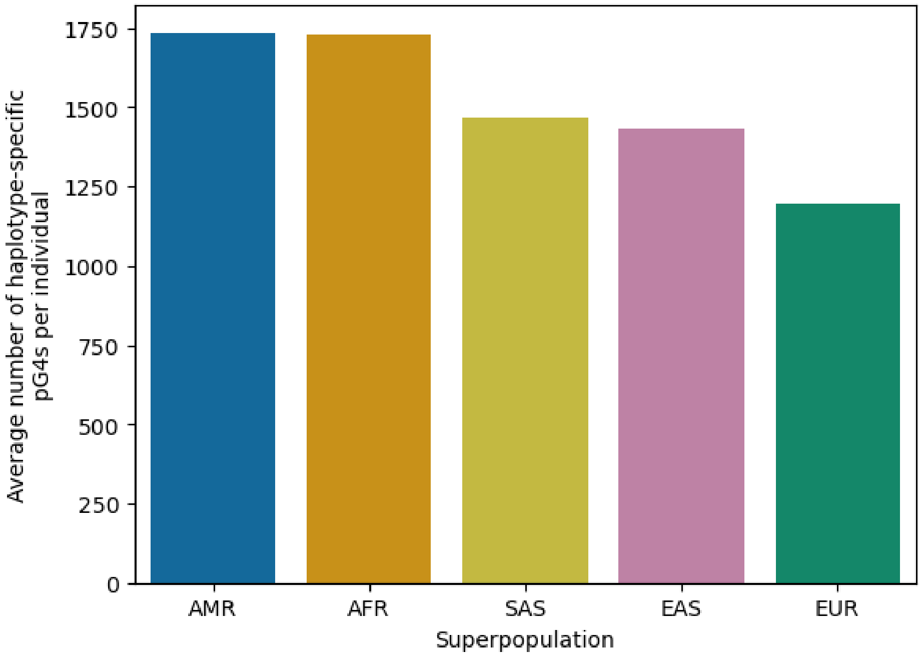

**Supplementary Figure 2**. The number of haplotype-specific pG4s in each sample across the pangenome.

1 and 2 represent the number assigned to each haplotype in the pangenome for any diploid sample. 0 represents the haploid assemblies.

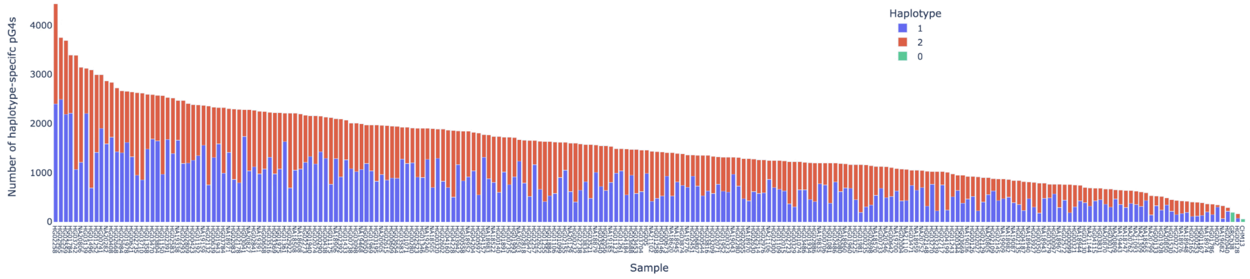

**Supplementary Figure 3**. The number of pG4s per pangenome sample that are absent from the reference assemblies (CHM13, GRCh38, and HG002).

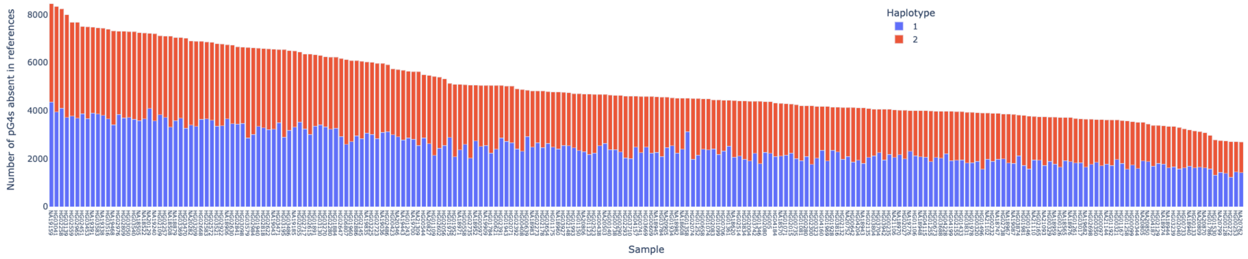

**Supplementary Figure 4**. Heatmap showing population sharing of pG4 families.

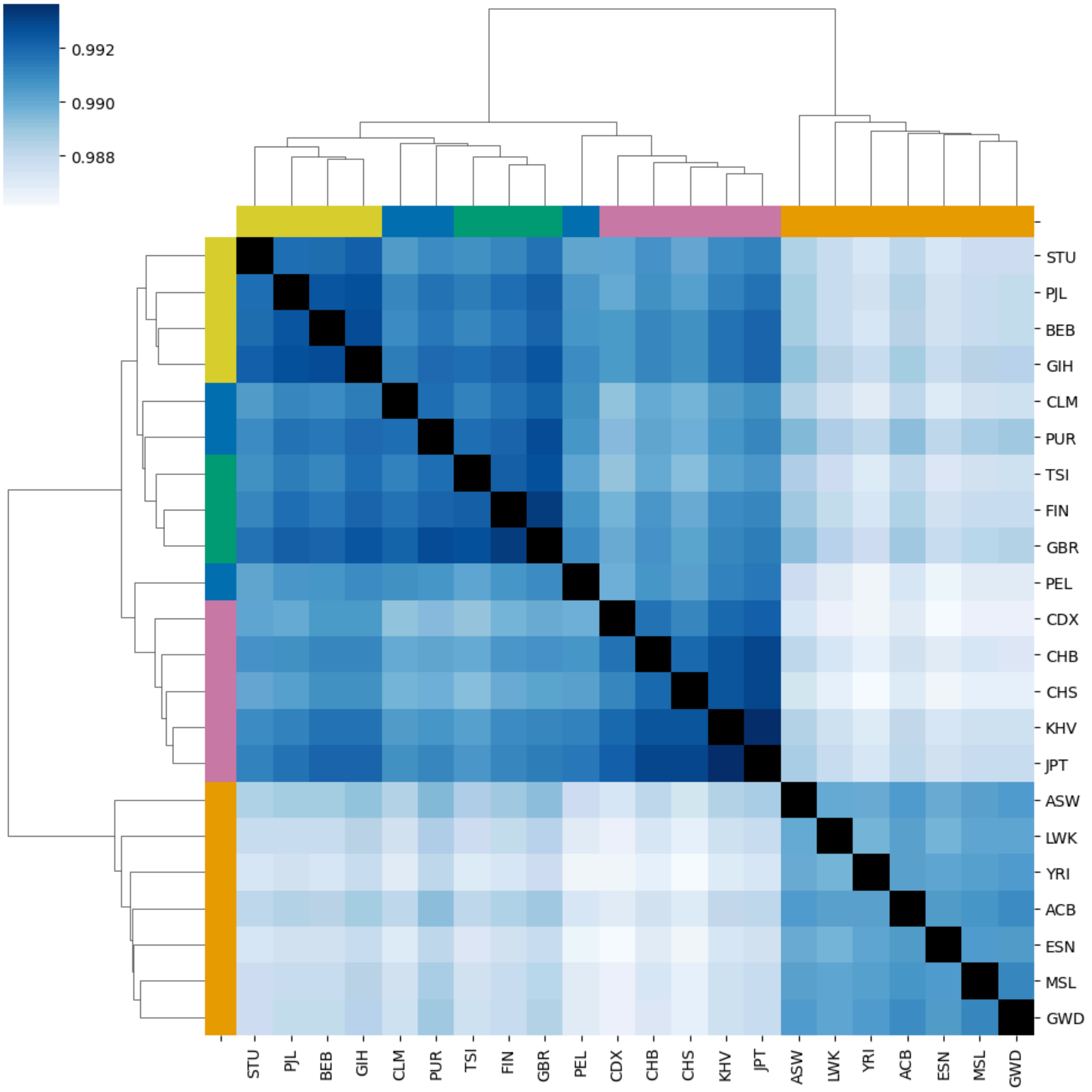

**Supplementary Figure 5**. The unrooted phylogeny, colored based on population labels.

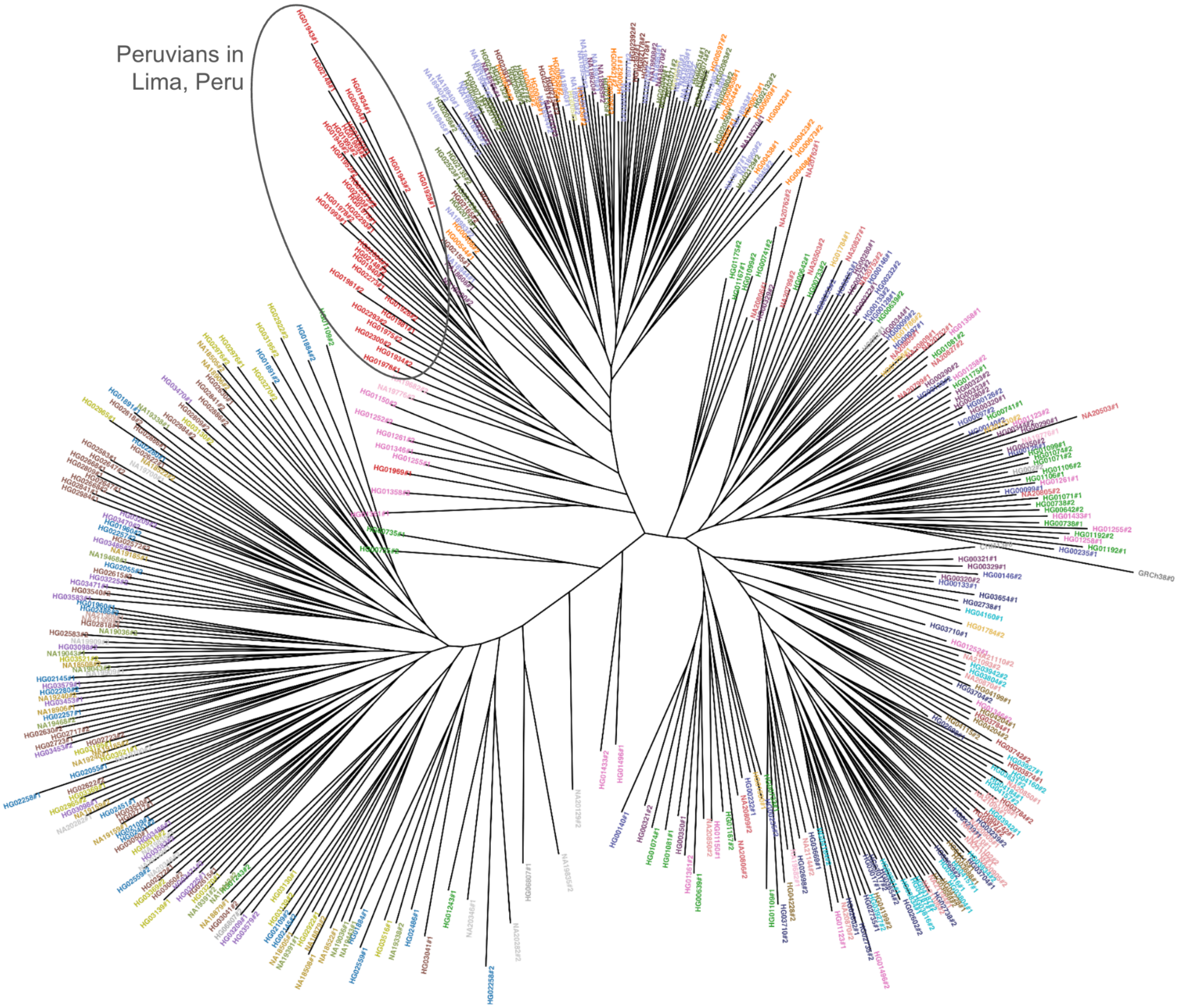

**Supplementary Figure 6**. pG4 families unique to populations.

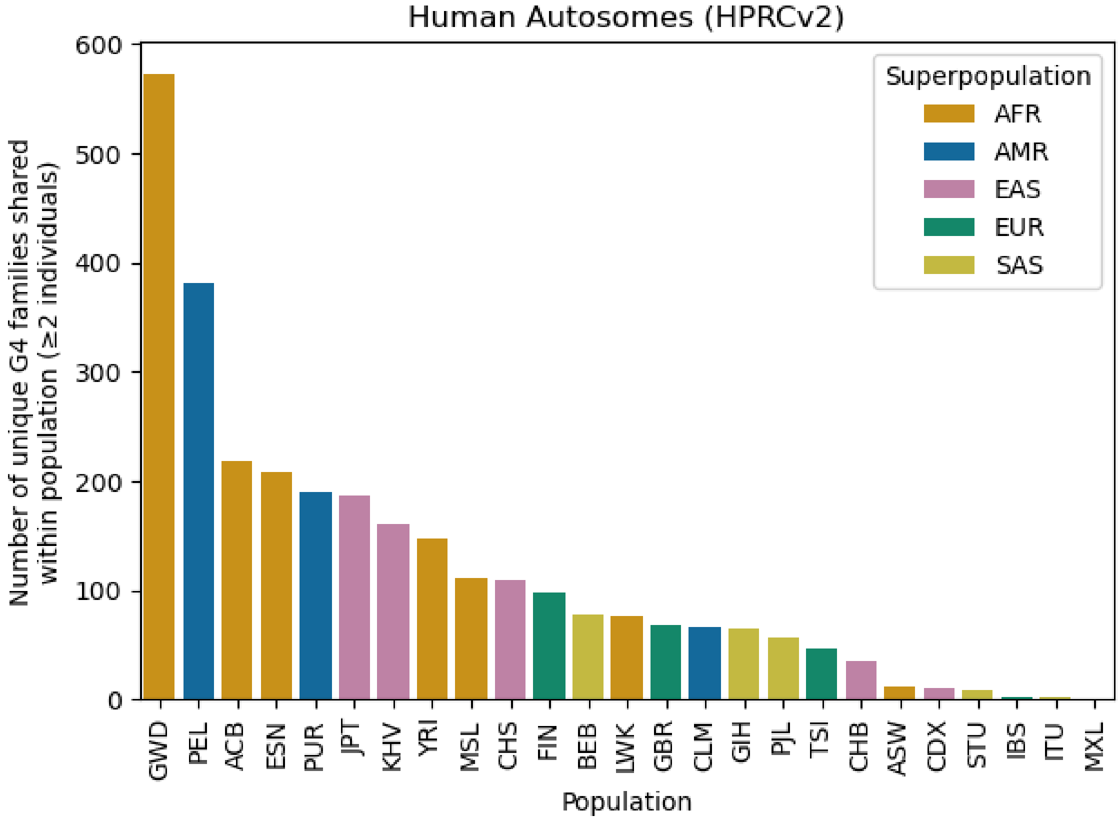

**Supplementary Figure 7.** PCA plot showing individuals of four populations clustering.

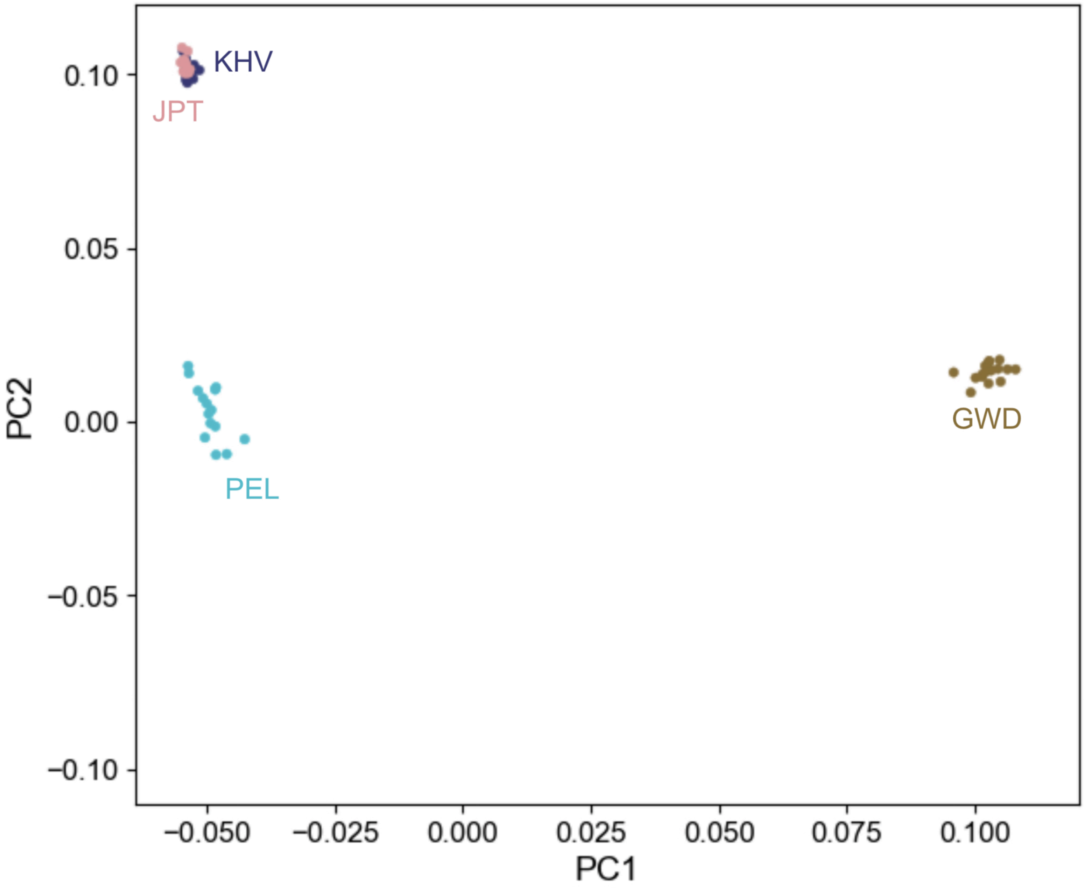

**Supplementary Figure 8**. Strand-specific pG4 selection coefficients across genic regions. Diamonds in light colors indicate regions where the model didn't fit the observed data with high confidence

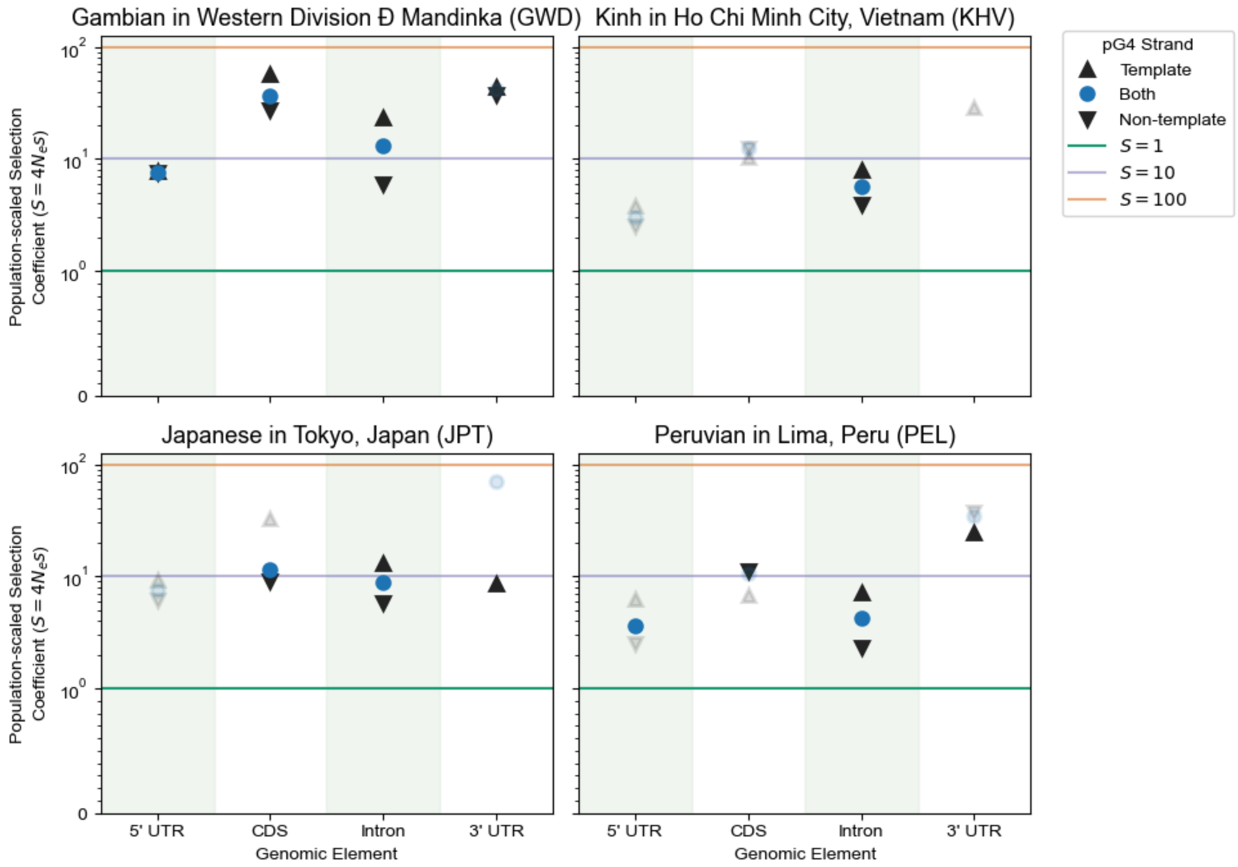

**Supplementary Figure 9**. Selection coefficients at all core, and stochastic origins of replications.

The non-confident values are not reported.

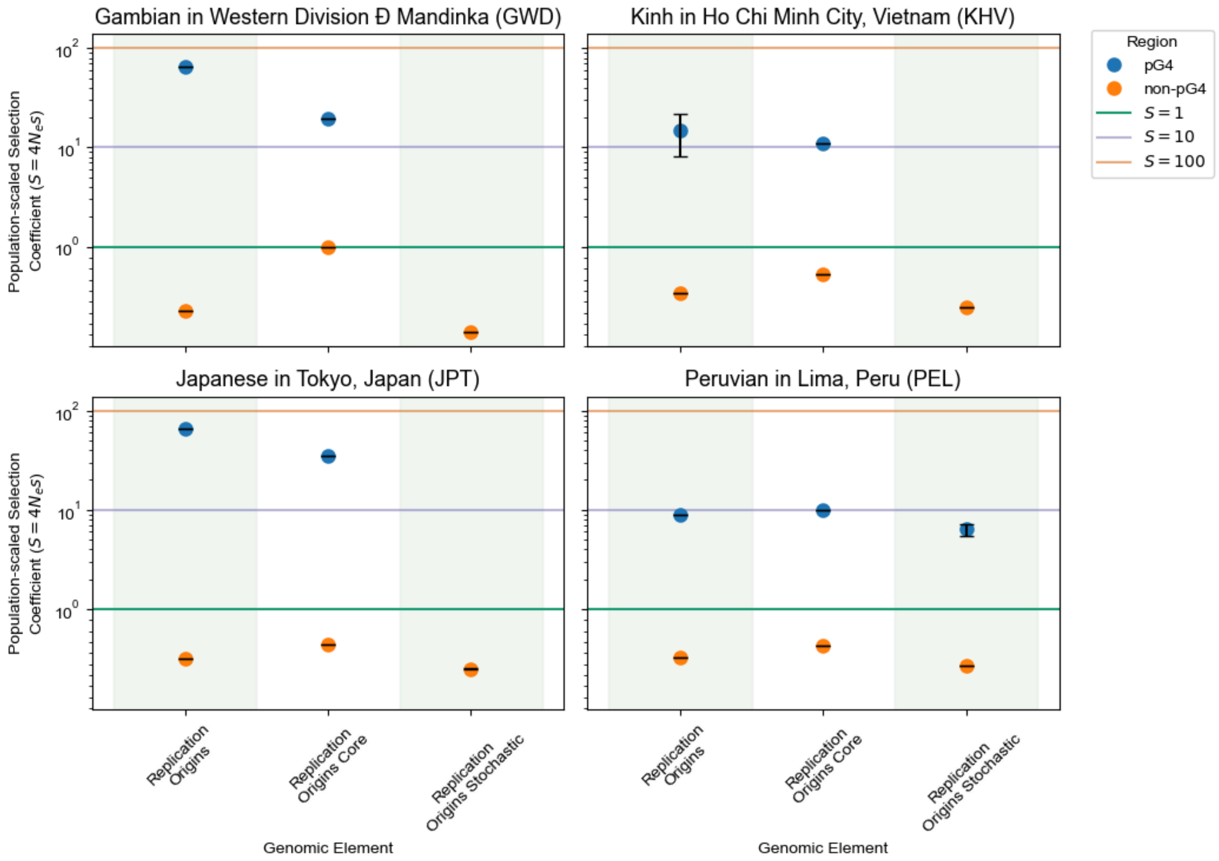

**Supplementary Figure 10**. Candidate genes with a significant pG4 variant associated with their expression.

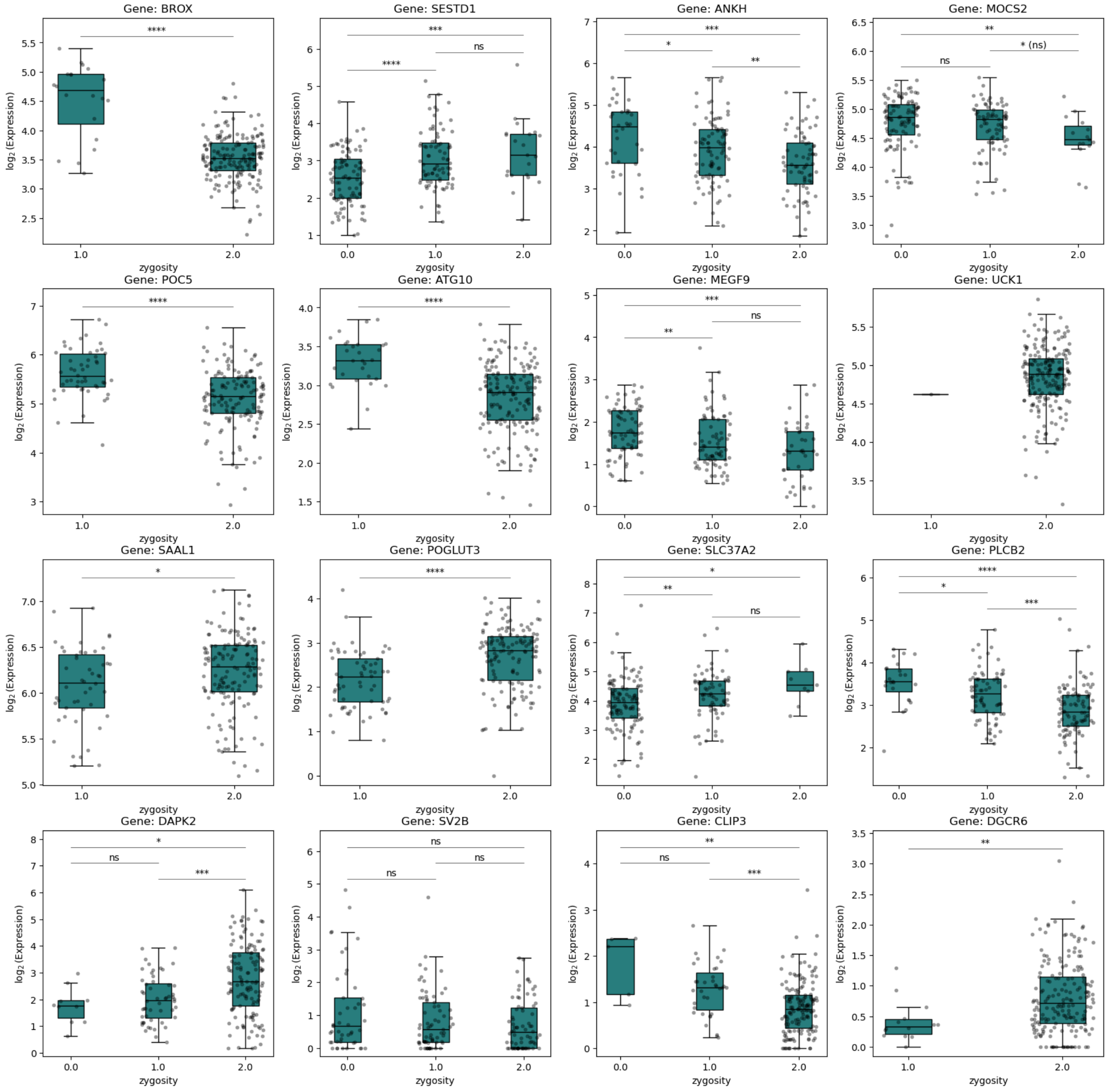

### **Supplementary Figure 11**. Additional candidate pG4s associated with alternative splicing.

We observed significant patterns of association with variants at pG4s at *FAH*, *TAF6*, *GIPR*, and *PROSER3* genes. The former two have pG4s on their template strand, while the latter two have pG4s on the non-template strand.

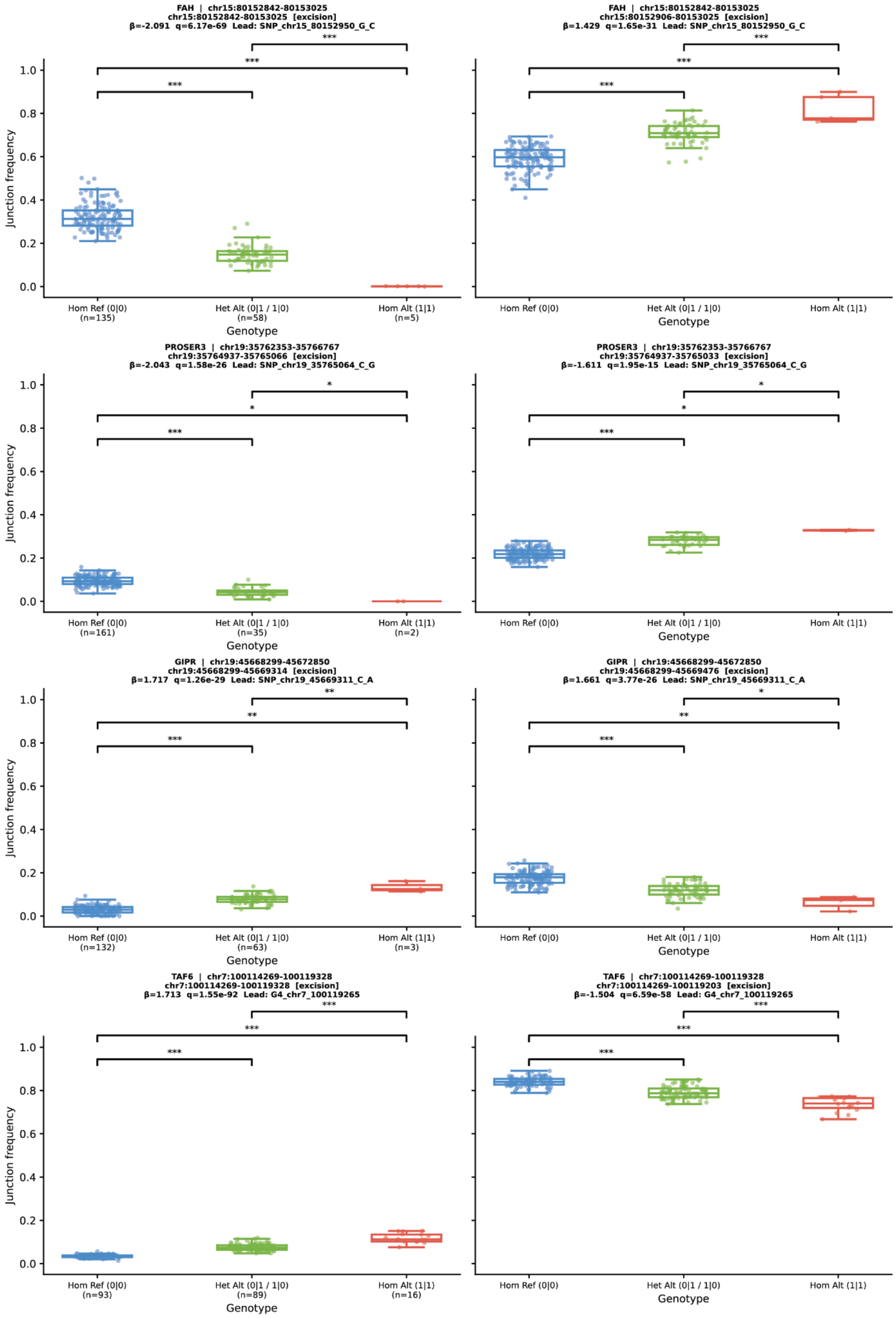

**Supplementary Figure 12**. The PAV matrix calculation schematic.

Highlighted columns show a pG4 family, and red dots are pG4s used in the analysis. Faded pG4s are removed from the analysis if they were either part of an unassigned or flagged region in any haplotype across the pangenome.

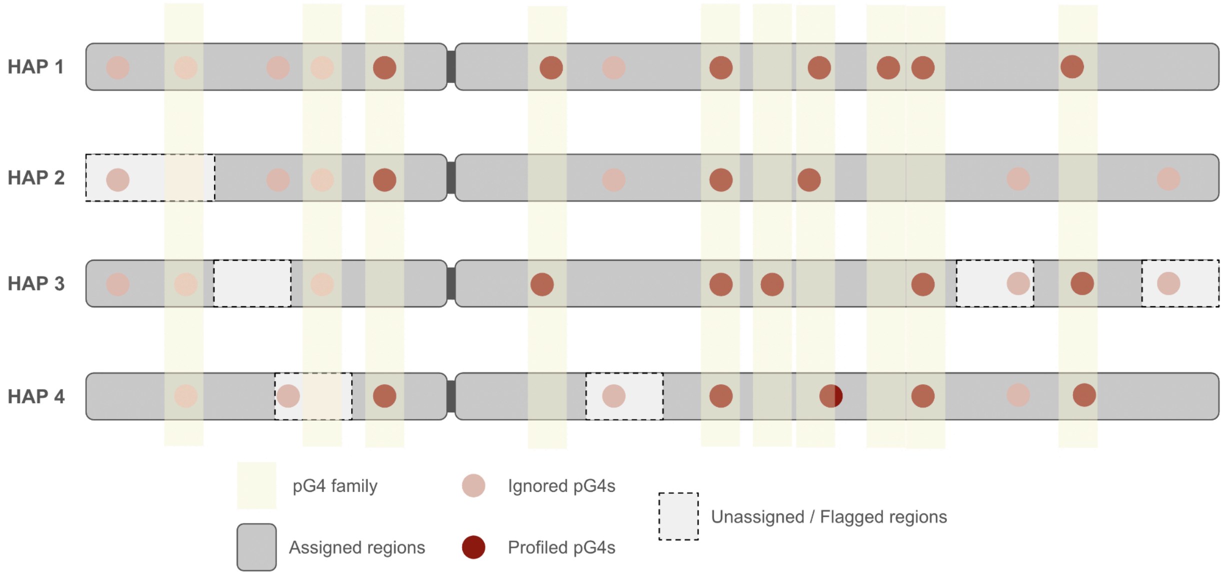
